# Increased expression of a subset of genes within reduced copy number regions across multiple cancer types

**DOI:** 10.64898/2026.04.10.717791

**Authors:** Tiffany A. Melhuish, Sara J. Adair, Anant Shah, Todd W. Bauer, David Wotton

## Abstract

The *TGIF1* transcription factor gene is present on chromosome 18, which is subject to whole chromosome copy number reduction in colon cancer. Despite this, *TGIF1* expression is significantly higher in cancer than in normal. In mice complete deletion of *Tgif1* reduced tumor burden in an *Apc* mutant model of intestinal cancer. Here we show that reducing *TGIF1* expression in a human colon cancer cell line slows proliferation and reduces growth of orthotopic xenografts. To ask if additional genes with copy number loss are more highly expressed in tumors we identified chromosomal regions subject to copy number reductions from ten TCGA cancer datasets. Within these regions a small proportion of genes, generally less than 10%, are expressed at higher levels in the tumor than in corresponding normal samples. Enrichment analysis using a set of 435 genes that have copy number reduction and increased expression identified mitosis as the most enriched gene set and FOXM1 and E2F family transcription factors as potential regulators. For mitotic genes, the average expression increase in tumor compared to normal is independent of copy number. In contrast, while DepMap common essential genes are generally more highly expressed in cancer than normal tissue, the relative increase in expression tracks well with copy number. Similarly, expression differences for gene sets such as S-phase, rRNA processing and DNA repair show increased expression in cancer versus normal, but changes also track with copy number. Thus, genes with increased expression despite copy number reduction may represent the output of key pro-tumorigenic transcriptional programs and could be potential therapeutic targets.

## INTRODUCTION

Multiple genetic changes drive cancer progression, including single nucleotide variants (SNVs), amplification or deletion of larger regions of the genome, and rearrangements that create novel fusions [1–3]. Alterations that affect a single gene, such as small deletions or insertions and single SNVs, can result in inactivation of tumor suppressors or activation of proto-oncogenes. Oncogene activation or tumor suppressor inactivation may be the key initiating events in many cancer types. Structural variants, such as amplifications or deletions of an entire chromosome or large chromosomal region, affect large numbers of genes, even though only one or a few genes within the region may provide the selective advantage to the cancer cell [2,4]. Classic examples of oncogene activation include activation of KRAS or BRAF by SNVs that result in single amino acid changes creating constitutively active variants [5,6]. SNVs frequently affect the p53 tumor suppressor resulting in inactivation or alteration of function [7]. In colon cancer, the APC tumor suppressor is frequently inactivated by a combination of nonsense or frame-shift mutations and loss of heterozygosity, and this is likely the most frequent initiating event in colon cancer [8,9]. Loss of APC function is often followed by activating mutations in KRAS and mutation of p53 [10]. In addition to altering expression of well characterized oncogenes and tumor suppressors, copy number changes can alter expression of a larger number of additional genes that can still contribute to tumor development, albeit with less impact than known cancer drivers.

Large deletions can affect entire chromosomes, resulting in reduced copy number of not only the tumor suppressor(s) that drive selection for the loss, but also of hundreds of other genes. For example, the most frequent genetic alteration in GBM is heterozygous loss of the entirety of chromosome 10, which is likely driven by the presence of the *PTEN* tumor suppressor [11]. In colon cancer there is frequent copy number reduction affecting chromosome 18 that harbors genes such as *SMAD4*, *SMAD2* and *DCC* with potential tumor suppressor function [12,13]. In addition to the well-known tumor suppressors that are likely the main drivers of selection for loss of such large chromosomal regions, it is also possible that reduced expression of other genes within the deleted region might favor cancer cell fitness. However, it is also likely that among the passenger genes that are co-deleted with known tumor suppressors there are those that might be required either for basic cellular functions or whose expression might be beneficial for the tumor cell specifically.

The hallmarks of cancer identify key functional capabilities of cancer cells and describe the complexity of both genetic and phenotypic changes in cancer [14,15]. In addition to promoting proliferation and suppressing death, these features include metabolic reprogramming, immune evasion and increased invasion and metastasis, among others. Thus, it is clear that a large number of gene expression changes occur during tumor progression, in addition to early alterations in oncogene and tumor suppressor expression or activity. Genome instability, and the resulting in increase in aneuploidy, are linked to tumor progression [16,17]. Increased aneuploidy can further promote tumorigenesis, by causing loss of heterozygosity for tumor suppressors, driving oncogene amplification, or by altering additional gene expression programs that can have cancer type specific effects [17].

TGIF1 was identified by its ability to bind to a specific retinoid response element, and can repress activation by retinoid X receptors [18]. TGIF1, and the closely related TGIF2 are also able to limit the transcriptional response to transforming growth factor (TGF) β signaling via interaction with TGFβ-activated SMAD transcription factors [19,20]. Given the tumor suppressive function of TGFβ signaling in many solid cancers [21], increased expression of TGIFs might be expected to promote tumorigenesis by limiting this pathway. In human gene expression data sets, expression of both *TGIF1* and *TGIF2* is significantly increased in colon cancer compared to normal tissue, and their expression is also increased in mouse colon tumors [22]. In an *Apc* mutant mouse model of intestinal cancer TGIFs promote tumorigenesis, although this appears to be by regulating gene expression via direct DNA binding independent of TGFβ signaling [22]. TGIF1 function has been implicated in the progression of additional cancer types. Increased TGIF1 expression promotes proliferation in esophageal cancer [23,24] and has been implicated in promoting lung adenocarcinoma and glioma [25,26]. In contrast, TGIF1 appears to be tumor suppressive in pancreatic ductal adenocarcinoma (PDAC). TGIF1 loss was shown to increase TGFβ signaling and drive EMT via a mechanism that may involve changes in both the expression and activity of TWIST1 [27]. Additionally TGIF1 loss was proposed to drive EMT by affecting both CD44 signaling and a PD-L1 immune checkpoint in PDAC [28]. Thus, TGIF1 may play context dependent roles that can be either tumor suppressive or pro-tumorigenic.

Despite the increased *TGIF1* expression seen in colon tumors compared to normal tissue, it has been suggested that TGIF1 loss of function may be a driver of colon cancer. Somatic mutations in *TGIF1* were identified in colon cancer in two studies, although in both the major effects on the *TGIF1* gene were reduced copy number and a paradoxical increase in expression [29,30]. The apparently contradictory changes in copy number and gene expression in colon cancer prompted us to further examine the role of TGIF1 in this cancer. More broadly, this contradiction made us ask whether there are additional genes that fit this pattern: Present in genomic regions that are subject to frequent copy number loss, but more highly expressed in cancer than normal tissue. Genes that are reduced in copy number due to selection for loss of tumor suppressor genes in the same chromosomal region, may themselves be beneficial for the tumor, with their upregulation may contributing to tumor progression. This class of genes might, therefore, include potentially actionable targets for cancer therapy.

Building from data supporting a pro-tumorigenic role for TGIF1 in colon cancer, despite the observed copy number reduction, we analyzed additional cancer data sets for chromosomal regions with frequently reduced copy number. We show that a small proportion of genes, generally less than 10%, within these chromosomal regions are more highly expressed in the cancer than corresponding normal samples. This gene set is enriched for mitotic genes and likely contains other genes that are important for tumor progression. Given the cost to the tumor cell of increasing expression of a gene with reduced copy number, this limited gene set may represent an enriched selection of genes that are more likely to be important for tumor progression than the bulk of genes that are more highly expressed in a cancer.

## RESULTS

### Frequent copy number loss of *TGIF1* in colorectal cancer

We have previously shown that deletion of the genes encoding Tgif transcription factors from intestinal epithelium slows the growth of early adenomas in mice with a heterozygous deletion of the *Apc* tumor suppressor gene [22]. Deletion of *Tgif1* alone had some effect on tumor size in the small intestine (SI), and the combination of deleting both *Tgif1* and *Tgif2* further reduced SI tumor growth and resulted in smaller adenomas in the colon. Pan-cancer analysis of copy number alterations (TCGA-PanCan; cBioPortal) shows a relatively consistent copy-number gain for *TGIF2* across multiple solid tumor types, with alteration in colorectal cancer (CRC) being the second most frequent (Fig 1A). This is consistent with a pro-tumorigenic role for *TGIF2* in CRC. In contrast, results with *TGIF1* were more complex, with most solid tumors showing a mixture of both loss and gain (Fig 1B). Two notable exceptions were CRC and TGCT (testicular germ cell tumors), in which most tumors had copy-number loss for *TGIF1*. Indeed, looking only at copy number loss for *TGIF1*, CRC was the second most frequently affected tumor type, with nearly 60% of cancers having loss of one copy of the gene (Fig S1A). Similar results were seen in two other colon cancer datasets, with co-occurrence of *TGIF1* loss and *TGIF2* gain across all three datasets (Fig S1B).

**Figure 1.**
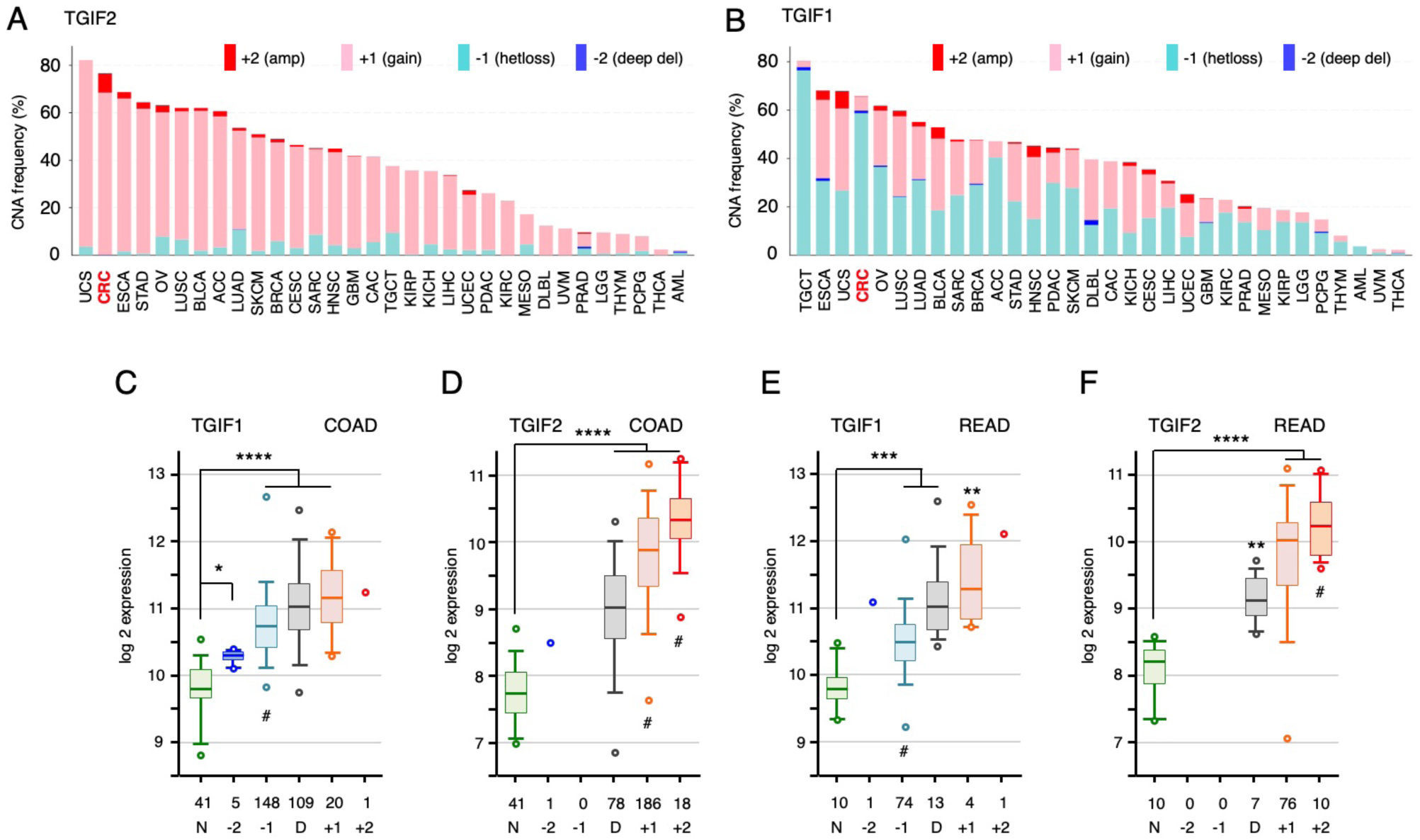
Copy number and expression analysis for TGIF genes. Copy number alterations for TGIF2 (A) and TGIF1 (B) are plotted across all TCGA PanCancer datasets (data from cBioPortal). C-F) Relative gene expression is plotted for TGIF1 (C) and TGIF2 (D) in the TCGA COAD dataset. Samples were separated into normal (N), and by predicted copy number among the tumor samples (-2: deep deletion, -1 heterozygous loss, D: diploid, +1: single copy gain, +2: gain of > 1 copy). The numbers for each are shown. The same analysis is plotted for the READ dataset (E, F). Plots (log2) are median, quartiles (box), 5^th^ and 95^th^ percentiles (whiskers), and extremes (circles). Adjusted p-values (student’s T-test): **** < 0.00001, *** < 0.0001, ** < 0.001, * < 0.01, compared to normal shown above for panels C-F. Asterisks in parentheses, below, indicate comparisons to diploid.

We next analyzed relative gene expression as a function of copy number for both *TGIF1* and *TGIF2*, this time analyzing colon and rectal cancers separately. As shown in Figure 1C, *TGIF1* expression was higher in the tumor samples than normal colon tissue, irrespective of copy number, with minimal difference between diploid samples and those that had either lost or gained one copy of *TGIF1*. As with *TGIF1*, *TGIF2* expression was significantly higher in tumors than normal, but appeared to scale with copy number, such that tumors with increased copy number had significantly higher expression than both normals and diploid tumor samples (Fig 1D). Similar results were observed for both genes in rectal cancer (Fig 1E, F). Given some of the limitations of our previous analysis of intestinal tumorigenesis – that the mouse model addressed only early adenomas, and the Villin-Cre driver was primarily expressed in small intestine – we wanted to test the effects of *TGIF1* and *TGIF2* in human colon cancer cells, which represent a more advanced form of cancer.

### Deletion of TGIFs slows proliferation

As a first test of the effects of TGIFs in human CRC, we mutated *TGIF1* and/or *TGIF2* using CRISPR/Cas9 in the human colon cancer cell line, HCT116. Following transfection with a CRISPR/Cas9 plasmid and a linear DNA fragment containing a selectable marker flanked by short homology arms, resistant colonies were selected and screened for integration of the resistance marker on one allele and an indel on the other. Indels were verified by sequencing PCR fragments, and loss of protein expression was tested by western blot (see Fig 2F,G for example). In addition to the TGIF genes, we also created control clones by targeting *THUMPD3*, and selected only for integration of the selectable marker. As shown in Fig 2A, homozygous loss of either *TGIF1* or *TGIF2* reduced cell number in a five day growth assay, with a further reduction in cell number in the double *TGIF1;TGIF2* mutant. In contrast, the control targeted cells were not different from the parental line. To control for clonal differences, we compared four independent *TGIF1;TGIF2* double mutants to four control clones for growth over 7 days and found significantly fewer cells at days 4 and 7 (Fig 2B) and reduced EdU incorporation, consistent with fewer cells entering S phase (Fig 2C). Plating cells at low density in a clonogenic assay demonstrated small decreases in colony number for either *TGIF1* or *TGIF2* mutation, with a further decrease in the double mutant (Fig 2D). Similar results were seen with growth in soft agar (Fig 2E), although mutation of *TGIF1* alone was not significant in this assay and there appeared to be greater combinatorial effect of the double mutant.

**Figure 2.**
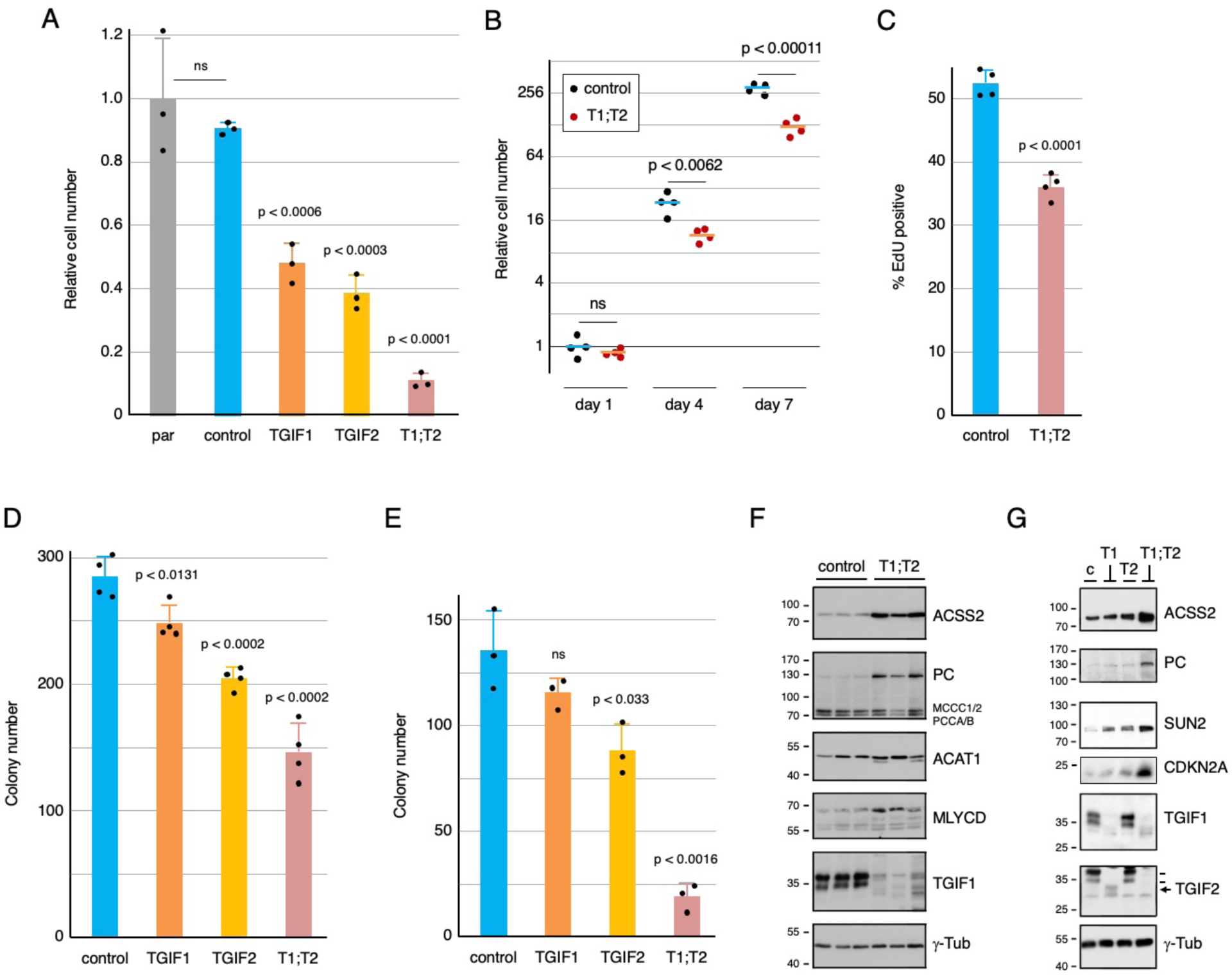
Inactivation of TGIF genes reduces HCT116 proliferation. A) Individual clones of HCT116 cells were analyzed for relative cell number. Triplicate cultures of each were counted on day 1 and day 5, and the relative cell number (mean +sd) was plotted relative to the parental cells (par). B) Four independent clones each for control and *TGIF1*:*TGIF2* null were counted at days 1, 4 and 7 and relative cell number (log2 scale) is plotted. C) % EdU positive cells is plotted for quadruplicate cultures for control and *TGIF1*:*TGIF2* null. D and E) Colony numbers for cells plated at low density on plastic (D) or in soft agar (E) are plotted. For A-E, p-values for comparison to the control cells are shown. Cell lysates were western blotted for the indicated proteins from three control and three *TGIF1*:*TGIF2* null clones (F) and from one clone of each of the four genotypes (the arrow indicates TGIF2, two lines show TGIF1 which is also detected by this antibody) (G). Molecular weight markers are shown to the left.

As in the mouse model, the above results suggest a combined effect of TGIF1 and TGIF2 on proliferation of human colon cancer cells. Analysis of RNA and protein from *Apc* mutant mouse colon tumors and from those also lacking both *Tgif* genes identified potential TGIF target genes [22]. We tested a small number of these by western blot in control and *TGIF1:TGIF2* double mutant clones, and found similar upregulation of ACSS2, PC (Pcx in mouse), ACAT1 and MLYCD in the double mutants (Fig 2F). Increases in expression were largely due to the combined loss of both TGIF1 and TGIF2, since more minimal effects were seen in either single mutant (Fig 2G). Taken together, these results suggest that complete loss of expression of TGIFs from a human colon cancer cell line has similar effects to deletion from mouse intestinal epithelium.

### Reduced TGIF expression slows tumor growth

To better model altered *TGIF1* or *TGIF2* expression rather than complete loss, we targeted *TGIF1* or *TGIF2* with validated shRNAs (Mission lentiviral shRNAs). As shown in Fig 3A, in three independent pools transduction of HCT116 cells resulted in a ∼60% reduction in *TGIF1* and ∼40% reduction in *TGIF2* expression compared to control cells. In a five day growth assay, this resulted in a small but significant reduction in cell number for either *TGIF1* or *TGIF2* knock-down (Fig 3B). To test the effects of reducing *TGIF1* or *TGIF2* expression *in vivo*, we generated orthotopic xenografts, by injecting human HCT116 cells into the wall of the cecum. To directly compare control and knock-down cells, we used an approach in which each cell type was differently tagged for identification by qPCR, combined and used to generate mixed tumors comprised of both control and knock-down cells in the cecum of *Foxn1* nude mice [31]. At necropsy, the relative proportion of each cell type was then determined by qPCR and normalized to that in the starting mix (see Fig 3C; [32]).

**Figure 3.**
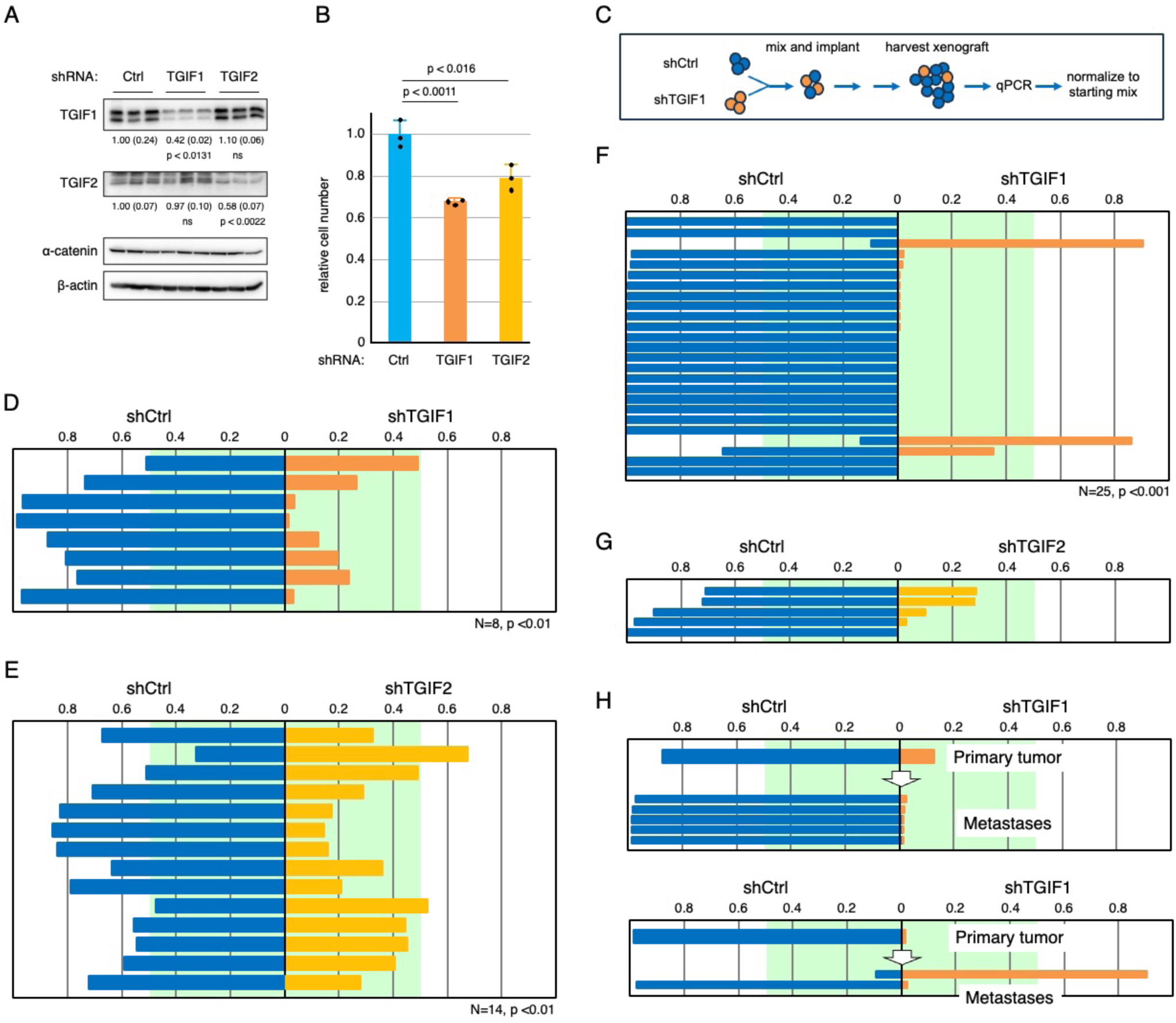
Reducing TGIF1 or TGIF2 expression slows proliferation. A) HCT116 cells were infected with lentiviral shRNA vectors targeting either TGIF1, TGIF2, or a control vector. Three independent pools were analyzed by western blot for TGIF1 and TGIF2. Expression relative to b-actin and a-catenin is shown. B) Triplicate cultures of control or TGIF1 or TGIF2 knock-down cells were analyzed for relative cell number after five days. Mean and sd are plotted (relative to cell number at plating). C) A schematic of the mixed xenograft approach used is shown. D-G) Comparison of relative proportions shCtrl and shTGIF1 cells in primary cecal tumors (D) and liver metastases (F), or the proportions of shCtrl and shTGIF2 cells in primary tumors (E) and liver metastases (G). Tumor number and p-value (Wilcoxon signed rank) are shown below each. H) Data from two mice are shown, indicating the proportions of shCtrl and shTGIF1 cells in the primary cecal tumor and in the liver metastases identified in that mouse. All plots show the relative proportion of shCtrl cells to the left (blue) and the proportion of shTGIF1 (orange) or shTGIF2 (yellow) to the right. The green area indicates the 0.5 boundary.

We first compared a mix of three *TGIF1* knock-down pools and three shCtrl pools, mixed at equal proportions. Following tumor harvest and analysis, the relative proportions of control and *TGIF1* knock-down cells were determined by qPCR, and the proportion contributed by either shCtrl or shTGIF1 compared by Wilcoxon signed rank test. As shown in Figure 3D, of the eight tumors from mice that were inoculated with a mix of *TGIF1* and control knock-down cells, the control cells dominated in all eight, with only one being close to an equal mix. We then performed a similar experiment comparing tumors with a mix of shCtrl and *shTGIF2* cells and again found that shCtrl cells predominated, with only two of 14 tumors having a majority of *shTGIF2* cells (Fig 3E). These results suggest that reducing expression of either *TGIF1* or *TGIF2* slows growth of human orthotopic CRC xenografts, consistent with our *in vitro* data. We also observed spontaneous metastasis to the liver, which is a primary site for CRC metastasis in humans. From the mice generated with a mix of *TGIF1* and control knock-down cells we isolated 25 individual liver metastases, and there was a highly significant enrichment for control cells in these metastases (Fig 3F). From the shTGIF2 mix, we only found five liver metastases, and although the control cells dominated in all five, this was not a large enough number to reach significance (Fig 3G). Comparing the proportion of control to *shTGIF1* cells in metastases to that in the primary tumor from the same mouse suggested that in most cases the metastases were similar to the primary tumor, with a further reduction in *shTGIF1* cells, although occasionally the proportion in a metastatic site was very different from the final proportion in the primary tumor (Fig 3H). Together, these results suggest that similar to the effects observed in early adenomas in genetic mouse models, the growth of more advanced human colon cancer cells as xenografts is impaired by reducing expression of *TGIF1* or *TGIF2*.

### A subset of chromosome 18 genes is upregulated in colon tumors

We analyzed copy number changes for all genes across the genome in colon and rectal cancer datasets from TCGA. A large portion of chromosome 20 was frequently amplified in both cancer types, consistent with both the increased copy number and increased expression for *TGIF2*. In contrast, the whole of chromosome 18 shows reduced copy number in both cancer types (Fig S2). We next asked if other genes on the recurrently deleted chromosome 18 were also upregulated in tumors compared to normal tissue or whether this was specific to *TGIF1*. As shown in Figure 4A, a relatively small number of genes on chromosome 18 are expressed at significantly higher levels in colon tumors, while the majority are downregulated. Similar results were seen with rectal cancer, although there are fewer normal samples in this dataset, so significance values were lower (data not shown). A total of 11 genes had increased expression (p-adj < 0.0001, log2FC > 1.0) in colon tumors compared to normal, and they appeared to be scattered along the length of chromosome 18 (Fig 4B). Relaxing the cut-off (p-adj < 0.01, log2FC > 0.4) included an additional 15 genes, which also span the majority of chromosome 18 rather than being located in a single cluster.

**Figure 4.**
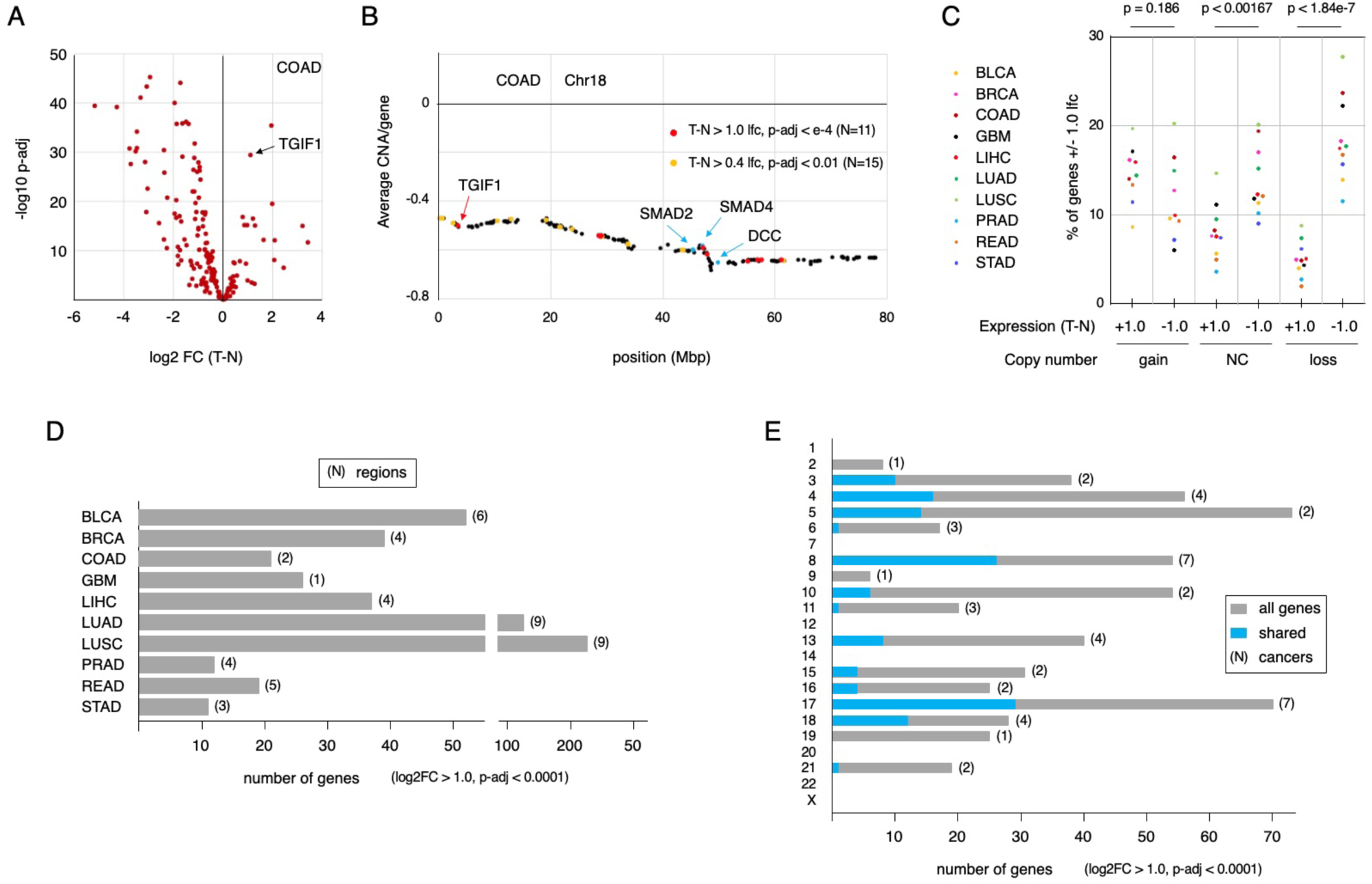
Analysis of copy number reductions in multiple cancers. A) A volcano plots for differential gene expression (tumor minus normal) for all genes on chromosome 18 is shown for the COAD dataset. TGIF1 is indicated by an arrow. B) Average copy number for all genes on chromosome 18 from the COAD dataset is plotted. The positions of TGIF1, SMAD2, SMAD4 and DCC are indicated, together with the positions of genes that have higher expression in tumor than normal (at two different stringency cut-offs). C) Gene expression data for each of ten TCGA datasets was divided by copy number (loss: < -0.4, gain: > +0.4, NC: -0.4 to +0.4), and the proportion of genes in each bin that were either increased or decreased (+/- 1.0 log2FC) in tumor versus normal was plotted for each. D) For each of the ten datasets analyzed, the total number of genes within recurrently deleted regions that have higher expression in tumor than normal (T-N log2FC > 1, p-adj < 0.0001) is plotted. The numbers in parentheses indicate the total number of regions with recurrently reduced copy number for each cancer. E) For each chromosome, the number of genes that have higher expression in tumor than normal (T-N log2FC > 1, p-adj < 0.0001) and are found in recurrently deleted regions is plotted. Total numbers are shown in gray, with the fraction that are shared between cancers shown in blue (for recurrently deleted regions that are found in 2 or more cancers). The numbers in parentheses indicate the numbers of cancers in which the recurrently deleted region is found.

### Multi-cancer analysis of gene expression and reduced copy number

*TGIF1* is one of a small number of genes on chromosome 18 with higher expression in CRC than normal tissue, and in mouse models, *TGIF1* promotes tumorigenesis. This prompted us to ask whether there are genes in other recurrently deleted regions of the genome that are overexpressed in cancer, and if so, whether they belong to specific gene sets. To expand our analysis of gene expression and copy number we examined data from ten TCGA PanCancer normalized datasets (UCSC Xena) and mapped chromosomal regions with altered copy number (Fig S2). For each data set we divided genes into one of three bins: increased (average CNA > +0.4) or decreased copy number (average CNA < -0.4), and minimal change (CNA between - 0.4 and +0.4). We then determined the proportion of genes within each bin that were significantly up- or downregulated in tumor compared to normal samples (log2FC > +1.0 or < - 1.0). There was a significantly larger proportion of downregulated genes than upregulated in the regions with minimal copy number change, and this difference was largely lost in regions that are amplified (Fig 4C). However, the most significant difference was between up- and downregulated genes in regions with reduced copy number, with relatively few being upregulated compared to normal. This is consistent with the idea that there is a cost to the cell to increase gene expression, and that this may be particularly evident for genes with reduced copy number. Thus, copy number reduced genes that are upregulated in the cancer may be upregulated because they are important for tumor progression or cell survival.

To identify genes that are present in larger copy number reduced regions but are upregulated in tumor compared to normal, regions with an average CNA of < -0.4 containing at least 100 genes (expressed in the relevant cancer or normal type) were included as a first selection (Fig S2, S3A). Our reasoning was that if there is selection for copy number reduction, presumably due to the presence of genes with tumor suppressive function, small deletions with very few additional genes are unlikely to be informative for this analysis. We also included some smaller deleted regions from other cancers, provided they reached the CNA cut-off and overlapped with one of the larger regions identified in another cancer type. For example, a 67 gene region in STAD was included since it overlapped with a 114 gene region from LUSC on chromosome 21 (Fig S3A). This analysis identified 47 regions with reduced copy number on 16 chromosomes, with at least two regions per cancer type found in nine of the ten cancers analyzed (Fig 4D, S3A). We next identified all genes with higher expression in tumor than normal (p-adj < 0.0001, log2FC > 1.0) across all 47 regions. For the majority of these regions, less than 10% of genes showed increased expression in tumor versus normal at this cut-off (Fig S3B). Since most regions with reduced copy number were shared between at least two cancers in this set, we compared the upregulated genes between the cancers. As shown in Figure 6C, the majority of genes with increased expression were unique to one cancer. This was true, even in cases such as the regions on chromosomes 8 and 17 (reduced copy number in 7 cancers each), where more than half of the upregulated genes were unique to a single cancer type (Fig 4E, Fig 5A,C). At the more stringent cut-off, no upregulated genes were shared between all seven cancers for the chromosome 8 and 17 regions (Fig 5A-D). Relaxing the fold change to > 0.4 (log2FC) and the adjusted p-value to < 0.01 increased the number of genes by about two-fold but did not greatly increase the number of shared genes among multiple cancers (Fig 5E, F). We also compared regions on chromosomes 5 and 10 that were shared between only two cancers each but contained relatively large numbers of upregulated genes. A significant fraction of the genes present in the reduced copy number region on chromosome 5 were upregulated in both BLCA and LUSC (Fig 5G). In contrast the overlap on chromosome 10 between GBM and LUSC was quite minimal (Fig 5H). We expanded this analysis using the less stringent expression cut-off for all pairwise combinations where at least 15 genes were upregulated (log2FC > 0.4, p-adj < 0.01) in both cancer types. The majority of these pairwise comparisons did not reveal significant overlap (Fig S4) or had too few genes that were upregulated within the reduced copy number region to reach significance. On chromosome 8, only LUAD and LUSC had significant overlap between upregulated genes, as did this pair for regions on chromosomes 3, 13 and 17. The other notable exception was the similarity between COAD and READ for genes in the chromosome 17 region (Fig S4B). Thus, while there are some recurrently upregulated genes within these reduced copy number regions, it appears that in general, this is not the case for most genes within these regions.

**Figure 5.**
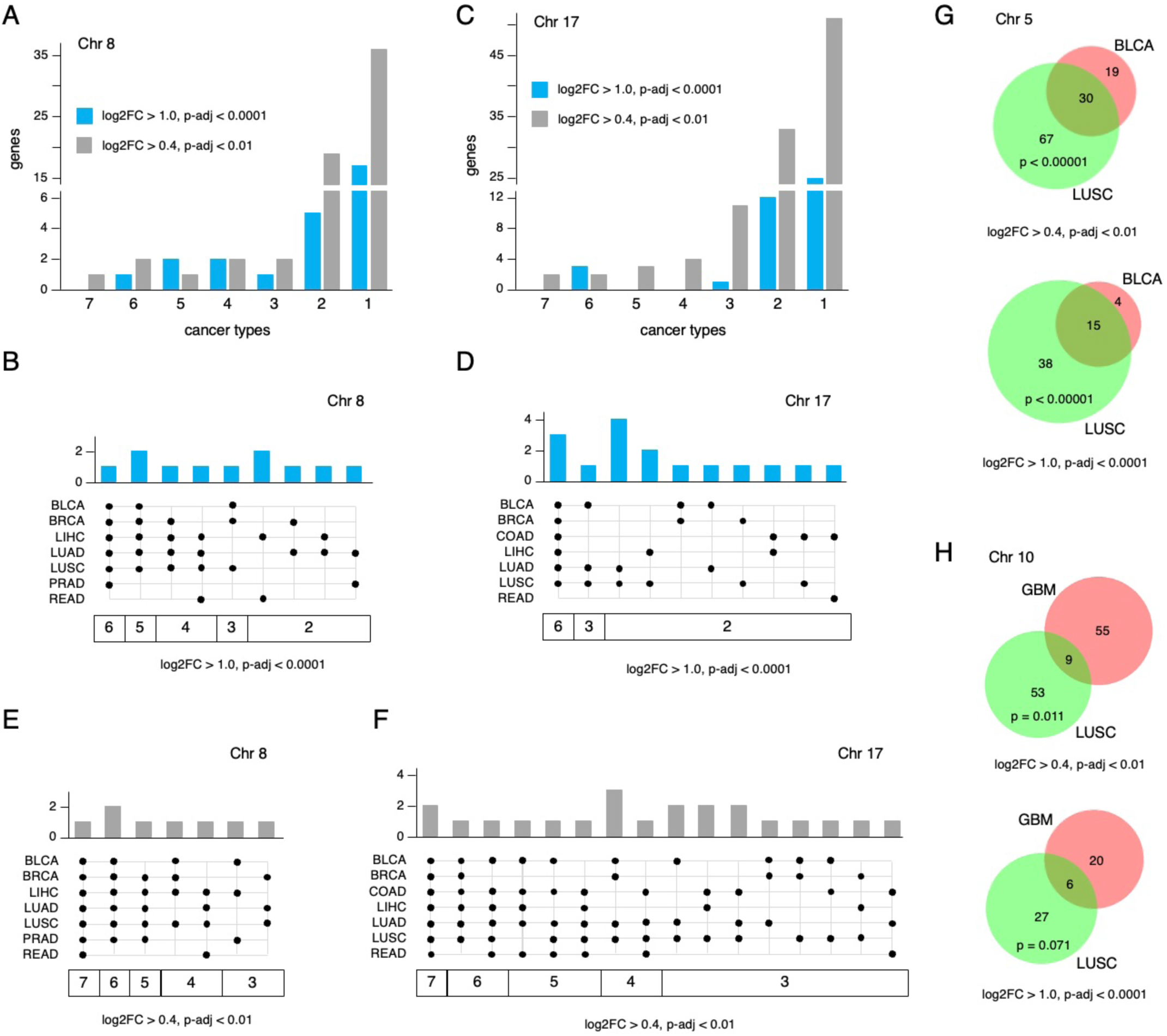
Increased gene expression in copy number reduced regions. The number of cancer types in which increased expression of all genes with higher expression in tumor than in normal is plotted for the copy number regions on chromosome 8 (A, B, E) and 17 (C, D, F). A and C show the number of genes upregulated (at two different stringency cut-offs) for one up to seven cancers. B and D show the cancers in which each recurrently upregulated gene is found, at the more stringent cut-off. E and F show the cancers in which each recurrently upregulated gene is found, at the more relaxed cut-off, for genes found in a minimum of three of seven cancers. G) The overlap between upregulated genes in a region on chromosome 5 that is found only in LUSC and BLCA is shown. H) The overlap between upregulated genes in a region on chromosome 10 found only in LUSC and GBM is shown. For G and H, overlaps are shown at two stringency cut-offs, and the p-value (hypergeometric test) for the overlap is shown.

**Figure 6.**
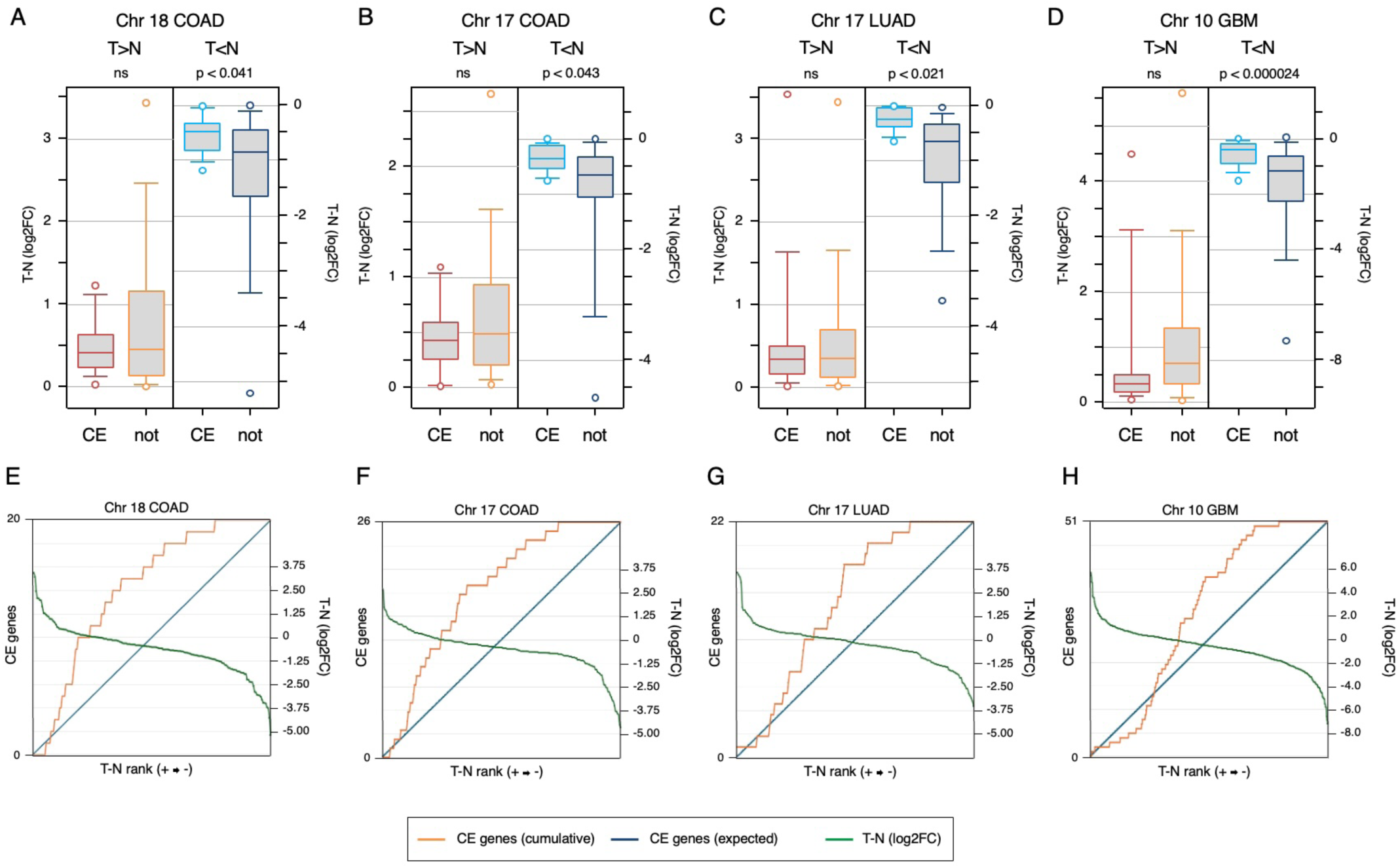
Common essential genes are less downregulated in copy number loss regions. A-D) For the four copy number reduced regions indicated the log2 fold change (tumor – normal) is plotted for four classes of genes: Common essential (abbreviated as CE) or not, as categorized by DepMap, and either higher (T>N) or lower (T<N) expression in tumor than normal. Plots are median, quartiles (box), 5^th^ and 95^th^ percentiles (whiskers), and extremes (circles). p-values shown are for comparison of CE vs not, for genes with either higher or lower expression in tumor than in normal. E-H) For the same regions shown in A-D, all genes within the copy number reduced region are ranked from highest log2 fold change (tumor – normal) to lowest and the position of each CE gene is plotted as cumulative function (orange; left hand axis), compared to the expected diagonal in blue. The log2 fold change for gene expression is plotted in green (right hand axis), with a log2FC = 0 centered on the 50^th^ percentile of the cumulative CE plot.

### Upregulated genes in commonly deleted regions are enriched for mitotic function

We next investigated what classes of genes are upregulated in regions with copy number reduction. As with the genes on chromosome 18 from COAD (see Fig 4B), upregulated genes in other chromosomal regions with reduced copy number did not appear to cluster based on chromosomal location (Fig S5). Possible interpretations of this include that the upregulated genes are indicative of common essential functions or represent the output of specific cancer-relevant transcriptional programs. Alternatively, it is also possible that this simply represents noise in the gene expression programs of these cancer types. To begin to address this, we compared the genes in some of the larger reduced copy number regions to the list of common essential genes from DepMap [33]. We first divided the genes within each region into four bins – higher or lower expression in tumor than normal, subdivided by common essential or not – and then compared the tumor versus normal log2FC among the four groups, with the expectation that, among the upregulated genes, common essential would be more upregulated. In contrast, we observed no statistical difference in tumor versus normal expression between upregulated common essentials and the remainder of the upregulated genes (red and orange; Fig 6A-D). In fact, although not significant, the trend was somewhat in the opposite direction. For genes with lower expression in tumor than normal there was a significantly smaller reduction in expression of the common essentials than the remainder of the downregulated genes (Fig 6A-D). Plotting the position of common essential genes across the expression change continuum from most upregulated in tumor to most downregulated supported the idea that the common essential genes within these regions were not those that change most between tumor and normal (Fig 6E-H).

Since this analysis suggests the upregulated genes are not predominantly classified as common essential, we next performed enrichment analysis on the list of 435 genes that were upregulated (p-adj < 0.0001, log2FC > 1.0) in at least one cancer type. GO term analysis revealed significant enrichment for mitosis and cell cycle (GO biological process) and for the mitotic spindle (GO cellular component) as the top hits (Fig 7A). Reactome pathways were also significantly enriched for cell cycle and mitosis. The most enriched class from ENCODE transcription factor ChIP-seq data was FOXM1, a key transcription factor for driving expression of mitotic genes (Fig 7A) [34]. To further examine the relationship between change in copy number and expression, we compared the average log2-fold change (tumor versus normal) between genes with copy number loss, gain, or no change (CNA > -0.4 and < 0.4). For this analysis we excluded PRAD and STAD since too few genes were affected by copy number alterations. As shown in Fig 7B, average differential expression between tumor and normal tracked with copy number – the log2-fold change was significantly lower with copy number loss than for genes with copy number gain. Interestingly, comparing the DepMap common essential genes showed a similar trend, but in all three bins the log2-fold difference between tumor and normal was greater, such that, even with reduced copy number, average gene expression did not decrease in the tumors (Fig 7B). In contrast to the results with common essential genes, mitotic genes were more highly expressed in tumor than normal independent of copy number change (Fig 7C). Surprisingly, other cancer related gene sets, including S-phase and DNA repair genes, did not show this same copy-number independence of gene expression change (Fig 7D-F, and Fig S6). Together, these data suggest that, across many chromosomal regions with reduced copy number in multiple solid tumor types, there is an enrichment for the upregulation of mitotic genes despite reduced copy number.

**Figure 7.**
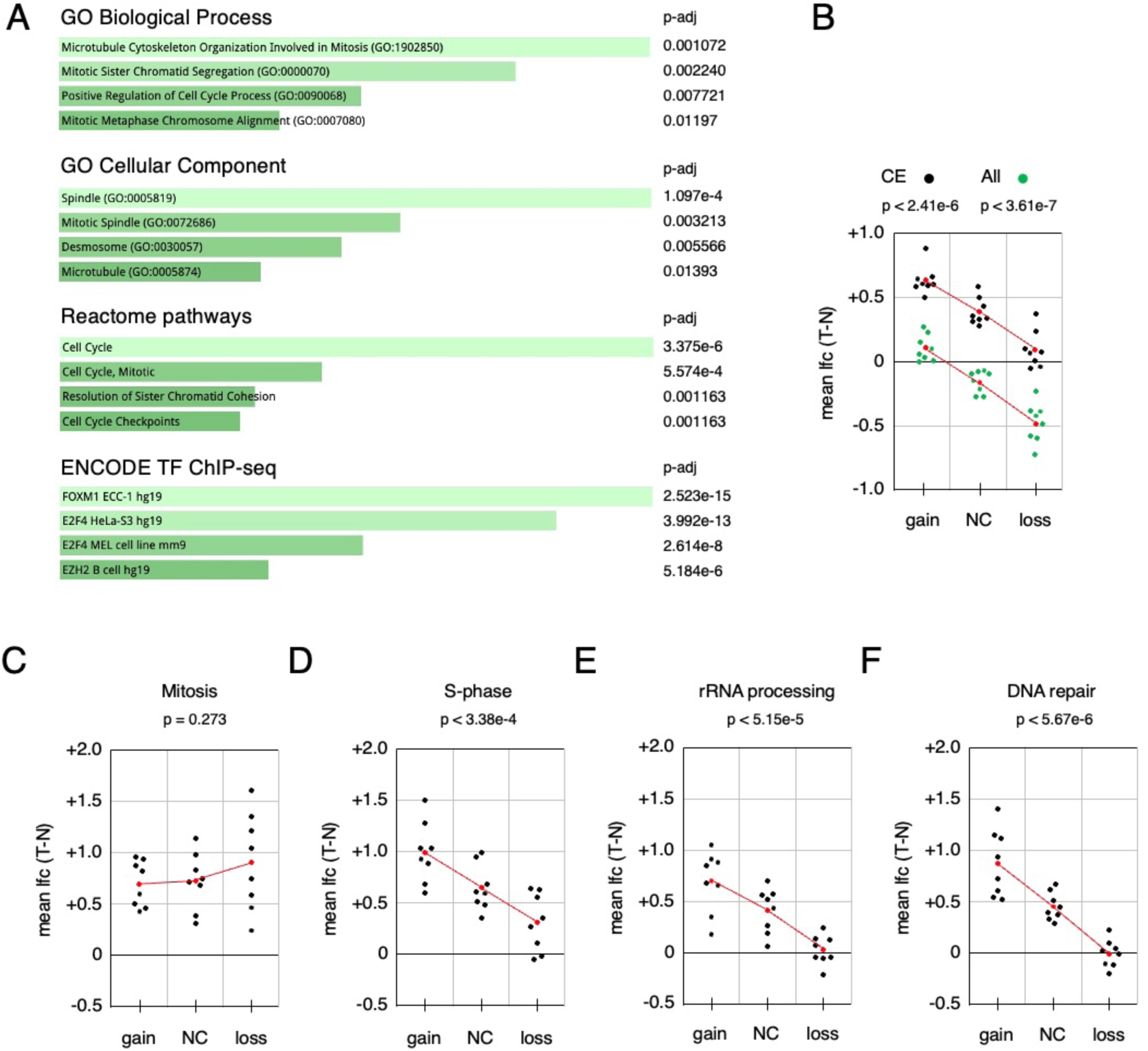
Enrichment analysis of genes in copy number reduced regions with higher expression. A) A total of 435 genes with higher expression in tumor than normal (T-N log2FC > 1, p-adj < 0.0001) in at least one region were subjected to enrichment analysis (Enrichr). The top four categories (ranked by adjusted p-value) are shown for the BP and CC GO terms, for Reactome pathways, and for transcription factor ChIP-seq from ENCODE. B-F) Using the same copy number bins as in Fig 6A, average tumor – normal log2 gene expression difference was plotted for each of the indicated gene sets. All genes and all common essential genes from DepMap were plotted together in B. p-values in B-F are for the comparison of average log2FC between the gain and loss categories. Plotted in red is the average for each bin, and the trendline based on the average.

## DISCUSSION

We have previously shown that deleting either *Tgif1* alone or both *Tgif1* and *Tgif2* together reduced the number of larger adenomas in a mouse model in which lesions are initiated by loss of the *Apc* tumor suppressor [22]. However, this model is limited, because the lesions do not progress beyond early adenomas. Additionally, due to the Cre driver this model primarily results in large numbers of adenomas in the small intestine, with relatively few colon tumors [22,35]. Although *TGIF1* and *TGIF2* are upregulated at the RNA level in human colon cancers, their function in human colon cancer cell lines or xenografts has not been addressed.

Consistent with the increased expression of *TGIF2* in human CRC compared to normal colon, the gene for *TGIF2* on chromosome 17 is subject to copy number gain in the majority of colon and rectal tumors. In contrast, TGIF1 is present on chromosome 18, which has reduced copy number in more than half of human colon and rectal cancers. Despite the reduced copy number, *TGIF1* expression is paradoxically higher in cancer than normal tissue when analyzing all tumors irrespective of copy number as well as when comparing only cancers with reduced copy number to normal samples. This suggests that specific mechanisms to increase expression of *TGIF1* in colon cancer are present that can compensate for the reduction in copy number.

The reduced *TGIF1* copy number seen in CRC is primarily a result of loss of a single copy of the entire chromosome 18. This is likely selected for in the cancer due to the loss of tumor suppressor genes such as *SMAD4*, which was originally identified as a tumor suppressor in pancreatic cancer (DPC4; [36]). SMAD4 is an essential mediator of TGFβ/SMAD signaling which is tumor suppressive in several cancer types, including colon [21,37], with inactivation of TGFβ signaling being one of the later steps of colon cancer progression [13,38]. In addition, SMAD2, which cooperates with SMAD4 to transduce TGFβ signals, is located on chromosome 18. The DCC (Deleted in Colon Cancer) tumor suppressor gene is also present on chromosome 18, and loss of both DCC and SMAD4 may contribute to colon cancer progression [12]. Reduced *TGIF1* copy number is likely a consequence of selection for loss of chromosome 18 tumor suppressors; however, cancer cells upregulate its expression. We previously demonstrated that increasing *TGIF1* expression in intestinal epithelium resulted in larger adenomas, at least in the small intestine [22]. In further support of this, we show that reducing (in contrast to the complete loss in the *Apc* mouse model) expression of either *TGIF1* or *TGIF2* in a human colon cancer cell line results in both a modest decrease in proliferation *in vitro* and a significant reduction in tumor growth in an orthotopic xenograft model. Importantly, there is a reduction in the fraction of *TGIF1* knock-down cells in both primary cecal tumors and spontaneous liver metastases, consistent with higher TGIF1 expression contributing to tumor growth. Interestingly, *TGIF1* point mutations have been identified in genome wide analyses of colon cancer, with the suggestion that these analyses identify mutations in driver genes [29,30]. However, the *TGIF1* mutations, some of which may be loss of function, have not been functionally validated, and are relatively rare. In this context, it is worth noting that in previous analyses examining missense variants in *TGIF1* in holoprosencephaly patients, we also identified close to half of them in the gnomAD database of unaffected individuals [39,40]. Thus, while we cannot rule out a pro-tumorigenic role for *TGIF1* loss of function mutations, in a subset of individuals or at a specific stage of cancer, it appears to be primarily increased *TGIF1* expression that contributes to colon cancer. Indeed, the main alterations in *TGIF1* in colon cancer are copy number reduction and increased expression [30].

Overall, our analysis supports a model in which increased TGIF1 expression contributes to tumor growth, despite the reduced copy number seen in a majority of colon and rectal tumors. This prompted us to ask whether other genes in chromosomal regions with recurrent copy number reductions are expressed at higher levels in tumors than in normal tissue. The rationale being that there is likely some cost to the cancer cell to increase expression of genes above that in normal tissue, particularly if those genes are present at reduced copy number. Consistent with this, we find that more genes in copy number reduced regions are downregulated in cancer than are upregulated, whereas this difference is not seen in amplified regions. For our initial analysis of copy number alteration and relative gene expression we examined the TCGA PanCancer COAD and READ datasets and included an additional eight PanCancer normalized datasets from TCGA, based primarily on whether the dataset included normal samples, with a preferential focus on those with larger numbers of samples. For these analyses we included only samples for which there were both copy number and gene expression data, but we did not sub-divide the expression data by copy number to avoid decreasing the numbers of samples in each sub-group. Additionally, genes with very low expression in both normal and cancer were excluded from the analysis, resulting in between 13989 and 14876 genes in each dataset. We identified large chromosomal regions with recurrent copy number reductions, most of which were not limited to a single cancer type. In most cases, where one of these regions was found in more than one cancer type the boundaries of the copy number alteration were quite similar between different cancers. For all regions, we identified a small number of genes with higher expression in tumor than in normal tissue, which in most cases represented less than 10% of the genes within the region with reduced copy number.

There are several possible explanations for why genes in regions with reduced copy number might be expressed at a higher level in tumors than in normal tissue. One trivial possibility is that this represents noise in the data, potentially due to multiple cell types present in the samples. Alternatively, the small fraction of genes that are more highly expressed could be a subset of genes not under strict transcriptional control in the tumors, so expression fluctuates stochastically, with some meeting our arbitrary cut-offs. Although we excluded genes with very low expression, this explanation may apply to the lower end of the expression range we did include. However, it is also possible for at least some of these genes that the higher expression is driven by the biology of the tumors and the function of these gene products. Our analysis of *TGIF1*, as an example, supports the idea that increased expression of at least some among these genes may contribute to tumorigenesis. More broadly, genes that encode essential functions might be expected to have increased expression irrespective of their copy number. Our comparison of genes with increased expression to the common essential gene list from DepMap argues against this, as common essential genes do not appear to be enriched among the most upregulated genes within copy number reduced regions. Common essential genes are generally upregulated in cancer compared to normal tissue, but as a group their relative expression appears to scale with copy number. However, gene set enrichment analysis suggests that there is a significant enrichment for mitotic genes among the genes with increased expression in tumor versus normal across all ten cancers analyzed. The most significantly enriched categories include functions related to microtubule organization, sister chromatid separation and the mitotic spindle, as well as the cell cycle more generally. Consistent with this, the most enriched ENCODE ChIP-seq transcription factor is FOXM1, which is a key driver of expression of mitotic genes [34]. In addition to FOXM1, MYBL2 and E2F family transcription factors contribute to mitotic gene expression, and enrichment for E2F target genes was also found in our analysis. FOXM1, and mitotic genes more broadly have been associated with increased copy number alterations and more aggressive cancers, and it is thought that the overexpression of mitotic genes, in part due to FOXM1 activity, contributes to aneuploidy [41,42]. Surprisingly, other cancer-relevant gene sets, such as S-phase or DNA repair genes do not show the same pattern as mitotic genes. In general, they have some increase in expression in tumor compared to normal, but as with common essential genes, the changes also track with copy number.

This enrichment for mitotic genes suggests that among the consistently upregulated genes present in regions of the genome that have reduced copy number there may be other non-mitosis related genes whose over-expression contributes to cancer progression. Our analysis of *TGIF1* is consistent with this idea: *TGIF1* expression is increased in CRC, it is present in a region of recurrent copy number loss, and reducing its expression reduces tumor growth in the intestine in both genetic and xenograft models. Given that there is likely a cost to the tumor of increasing expression of genes that are present at reduced copy number, focusing on such genes might be a simple way to enrich for those that are pro-tumorigenic from among the large number of gene expression differences between tumor and normal. It would, therefore, be of interest to functionally test other genes that fit these criteria, particularly those that are found upregulated in multiple cancers.

Mitosis was the only pathway that was clearly enriched in the minimal gene set identified as increased expression with copy number reduction. Where copy number reduced regions were shared between cancer types, individual mitotic genes tended to be found upregulated in multiple cancer data sets. However, the majority of genes identified in this manner were not shared between cancer types, even when the deleted region was present in several cancers. This suggests that there may be cancer-specific requirements for increased expression of other genes in this class. Our approach generated a relatively small gene set, which could be refined by comparison to gene products with existing small molecule inhibitors to further narrow down potential therapeutic targets for specific cancer types. In summary, we suggest that this approach has the potential to identify a small subset of all genes that are upregulated in a cancer type, that may include actionable targets.

## MATERIALS AND METHODS

### Cell culture and CRISPR/Cas9 deletion and shRNA knock-down

HCT116 cells were cultured in RPMI (Invitrogen) with 10% Fetal Bovine Serum (Hyclone). HEK293T were cultured in DMEM (Invitrogen) with 10% Fetal Bovine Serum (Hyclone) and were transfected with PEI (Sigma). HCT116 identity was verified by STR profiling. shRNA knockdown used validated Mission lentiviral shRNA vectors (Sigma; TRCN0000020153 - TGIF1, TRCN0000273612 - TGIF2). HCT116 cells were transduced with lentiviral vectors under standard conditions using polybrene (8ug/ml), and 24 hours after infection, cells were subjected to selection with 0.15μg/ml puromycin. After 7 days in selection, pools were frozen for xenograft generation and checked for knock-down by western blot. CRISPR/Cas9 mutants were generated by transient transfection in HCT116 cells using PEI. A Cas9 plus gRNA plasmid (pX330; Addgene 158973) was co-transfected with a PCR-generated linear DNA fragment that included a selectable marker (puromycin or zeocin) with ∼25bp flanking arms homologous to regions either side of the gRNA target. 48h after transfection, cells were re-plated at low density and subjected to selection with puromycin or zeocin. Resistant colonies were expanded and tested by western blot and Sanger sequencing of PCR fragments spanning the gRNA target sequence. pX330-gRNA was a gift from Charles P. Lai (Addgene plasmid # 158973; http://n2t.net/addgene:158973) [43].

### Antibodies and western blotting

Cells were lysed directly in SDS-PAGE loading buffer and boiled or were sonicated using a Branson digital sonifier with microtip in 1% NP-40 in PBS. Lysates were separated by SDS-PAGE, transferred to Immobilon-P (Millipore) and proteins visualized using ECL (Millipore, WBKLS0100). Primary antibodies were against ACSS2 (Cell Signaling #3658), TGIF1 [19], γ-tubulin (Sigma T6557), ACAT1 (Proteintech 16215-1-AP), MLYCD (Proteintech 15265-1-AP), SUN2 (Abcam ab124916), CDKN2A (Abcam ab80), TGIF2 (Proteintech 67576-1-Ig), β-actin (Proteintech 66009-1-Ig) and α-catenin (Epitomics 2028). PC was detected using Neutravidin conjugated HRP (ThermoFisher). Signals were quantified on a BIO-RAD ChemiDoc MP Imaging System and BIO-RAD Image Lab software.

### Proliferation and colony assays

Proliferation assays were carried out either by counting cells after 1, 3 and 7 days in culture, or by crystal violet assay: Cells in 12 well plates were washed with PBS, stained with crystal violet (0.2% crystal violet [Sigma, C6158], 2% ethanol in dH_2_O) for 10 minutes and washed twice with water. then The stained cells were solubilized in 1% SDS, and the absorbance measured at 570 nm. For EdU labeling, cells were seeded in 4-well chamber slides and treated as described. Cells were pulse labeled with 10μM EdU for 45 minutes, then fixed with 4% paraformaldehyde (PFA). EdU was detected using the Click-iT EdU Kit (Invitrogen C10339) following the manufacturer’s protocol and imaged as described [44]. Colonies in clonogenic assays in 6 well plates were stained with crystal violet as described above, then dried and imaged on a BIO-RAD ChemiDoc MP Imaging System. For growth in soft agar, cells were plated at 1000-5000 cells per well in 0.3% final agarose overlayed on a 0.5% agarose layer in 6 well culture dishes, grown for 3-4 weeks, stained with crystal violet, and imaged as above.

### Orthotopic xenografts

All animal procedures were approved by the Animal Care and Use Committee of the University of Virginia, which is fully accredited by the AAALAC. HCT116 cells with pLX304-PCR-tags and Mission shRNA constructs introduced by lentiviral infection were collected as pools within 2 weeks of introducing the shRNA. Cells were mixed at equal proportions one day prior to xenograft generation and cultured together overnight. Cells were collected by trypsinization and resuspended in antibiotic-free media at 16.67x10^6^/ml. Eight week old athymic nude mice (Jackson 007850) were injected with a total of 1 x 10^6^ cells in the cecal wall at six sites (10ul each) using a Hamilton syringe with three injections per side [31]. Tumors were allowed to progress until humane endpoints were reached, and then frozen prior to homogenization using Polytron homogenizer at 300 mg/ml in PBS. Genomic DNA was prepared by HotSHOT [45], and normalized by qPCR using 1ul for qPCR using common primers. Following normalization and 12 cycles of pre-amplification with common primers, qPCR was performed using 2% of the pre-amplified product (SensiMix SYBR + FITC PCR Mix [Bioline], myIQ thermocycler [BIO-RAD]). Proportions of each PCR-Tag were calculated by 2^ΔCt^ using the time zero mixture of cells as the reference [32].

### Datasets and analysis of expression versus copy number

CNA plots across all TCGA PanCancer datasets were generated in cBioPortal [46]. Gene set enrichment analyses were performed using Enrichr (https://maayanlab.cloud/Enrichr/) [47,48]. For all other copy number and expression analyses, TCGA PanCancer datasets (gistic2 thresholded CNA and IlluminaHiSeq expression data) were downloaded from UCSC Xena (https://xena.ucsc.edu/) [49]. Datasets were trimmed to include only samples present in both, and to remove normal samples (used for generating expression comparison to tumor samples). Gene lists were trimmed to remove genes with average expression that was not above 2 (log2) in either tumor or normal and were aligned with gencode hg19 probemap for chromosomal location. For CNA, average copy number across all samples for a particular cancer was plotted per gene, and relative gene expression was analyzed by comparing tumor to normal, including all tumor samples with CNA data, irrespective of copy number. Chromosomal regions with reduced copy number were initially selected if they contained at least 100 genes and average CNA was -0.4 or less. In additional cancers, regions with less than 100 genes with CNA < -0.4 were included if they fully overlapped with a region in another cancer that satisfied the primary cut-off. Additionally, if the majority of a reduced copy number region was CNA < -0.4 and was not the primary selector of this region, interspersed sub-regions with a CNA > -0.4, but less than -0.35 were tolerated. For the analyses comparing expression to copy number change genome-wide (Figs 5A,B, 9B-F) all genes meeting the expression level cut-off were separated based on copy number change and the proportion with a T-N log2FC >+/- 1 determined for each dataset or the average T-N log2FC for a specific gene set determined. For comparison of common essential gene expression (Fig 6), the average gene expression (T-N log2FC) was compared (student’s T-test) for genes within the indicated copy number reduced regions. Average gene expression (Fig 7B-F) was compared (student’s T-test) for specific gene sets between all genes with copy number gain or loss genome wide. Comparison to common essential genes used a list of CRISPR based common essentials from DepMap [33].

## ACKNOWLEDGEMENTS

This work was supported by funding from the NCI: R01CA259571, and by the NCI Cancer Center Support Grant (P30 CA44579) to the University of Virginia Comprehensive Cancer Center. The authors thank the Research Histology Core and Molmart for assistance, and members of the Center for Systems Analysis of Stress-Adapted Cancer Organelles for helpful discussion and advice.

## Figure Legends

**Figure S1.**
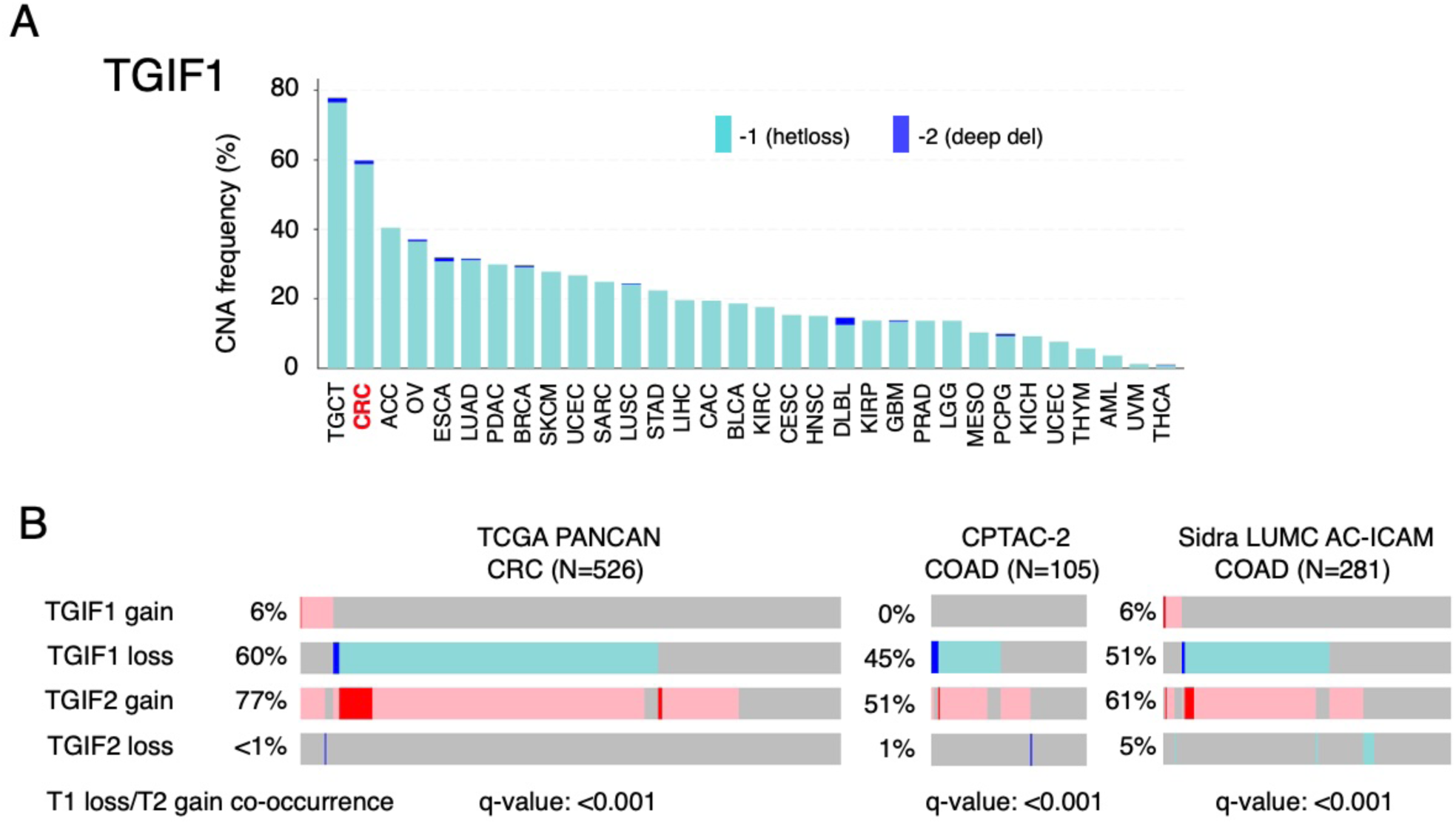
TGIF1 copy number reductions in PanCancer datasets. A) Copy number reductions for TGIF1 are plotted across all TCGA PanCancer datasets (data from cBioPortal). B) Comparison of CNA (loss and gain) for TGIF1 and TGIF2 in the TCGA PanCancer CRC dataset and in two additional COAD datasets from cBioPortal. q-values for co-occurrence of TGIF1 CNA reduction and TGIF2 CNA gain are shown below.

**Figure S2.**
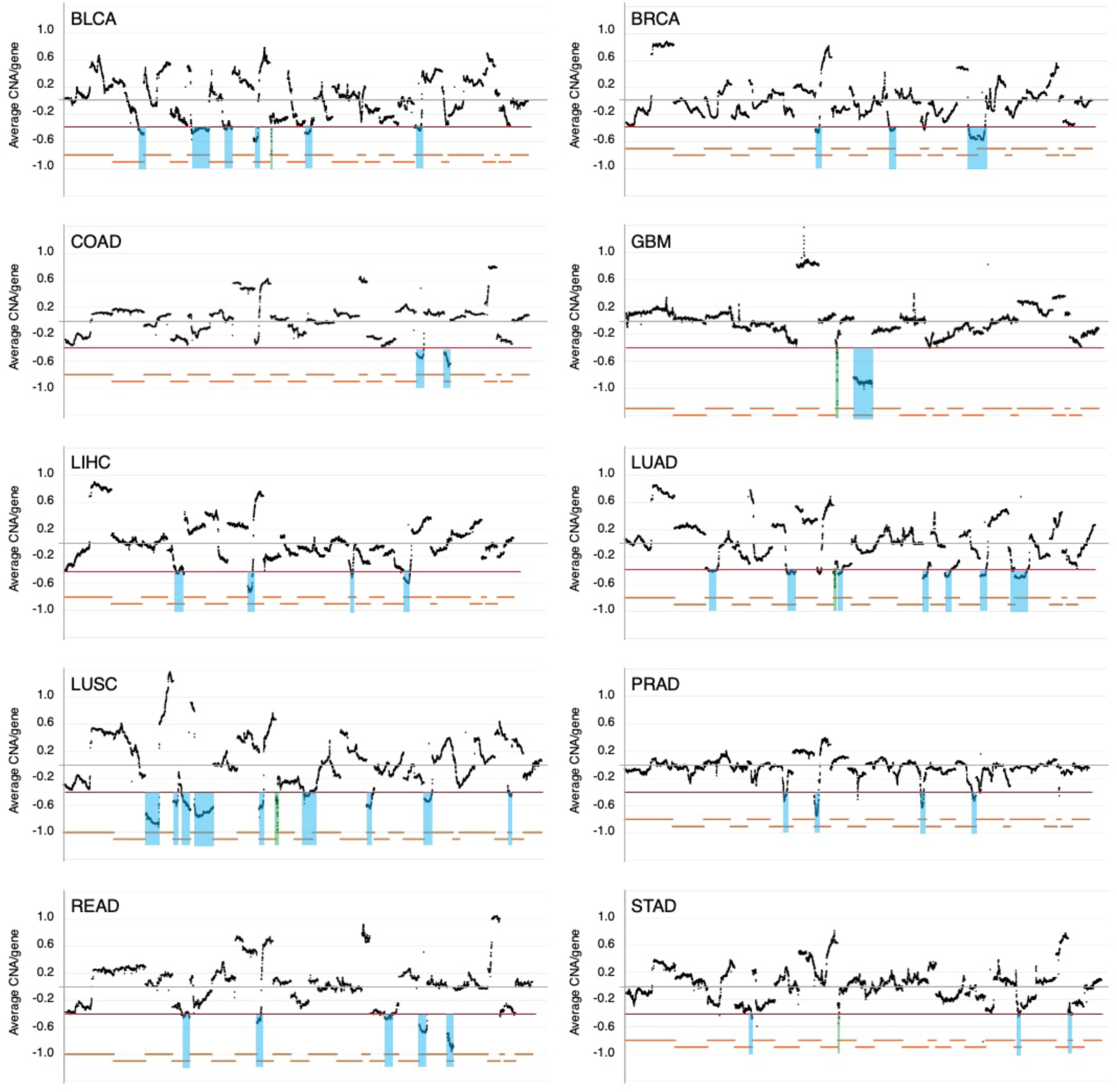
Analysis of recurrent copy number in TGCA PanCancer datasets. Average copy number per gene is plotted genome wide for ten TCGA PanCancer datasets. Red lines indicate the average CNA -0.4 cut-off used to select regions, which are indicated in blue. Chromosomes are ordered 1 to 23 (left to right) and are indicated by the orange lines below each plot.

**Figure S3.**
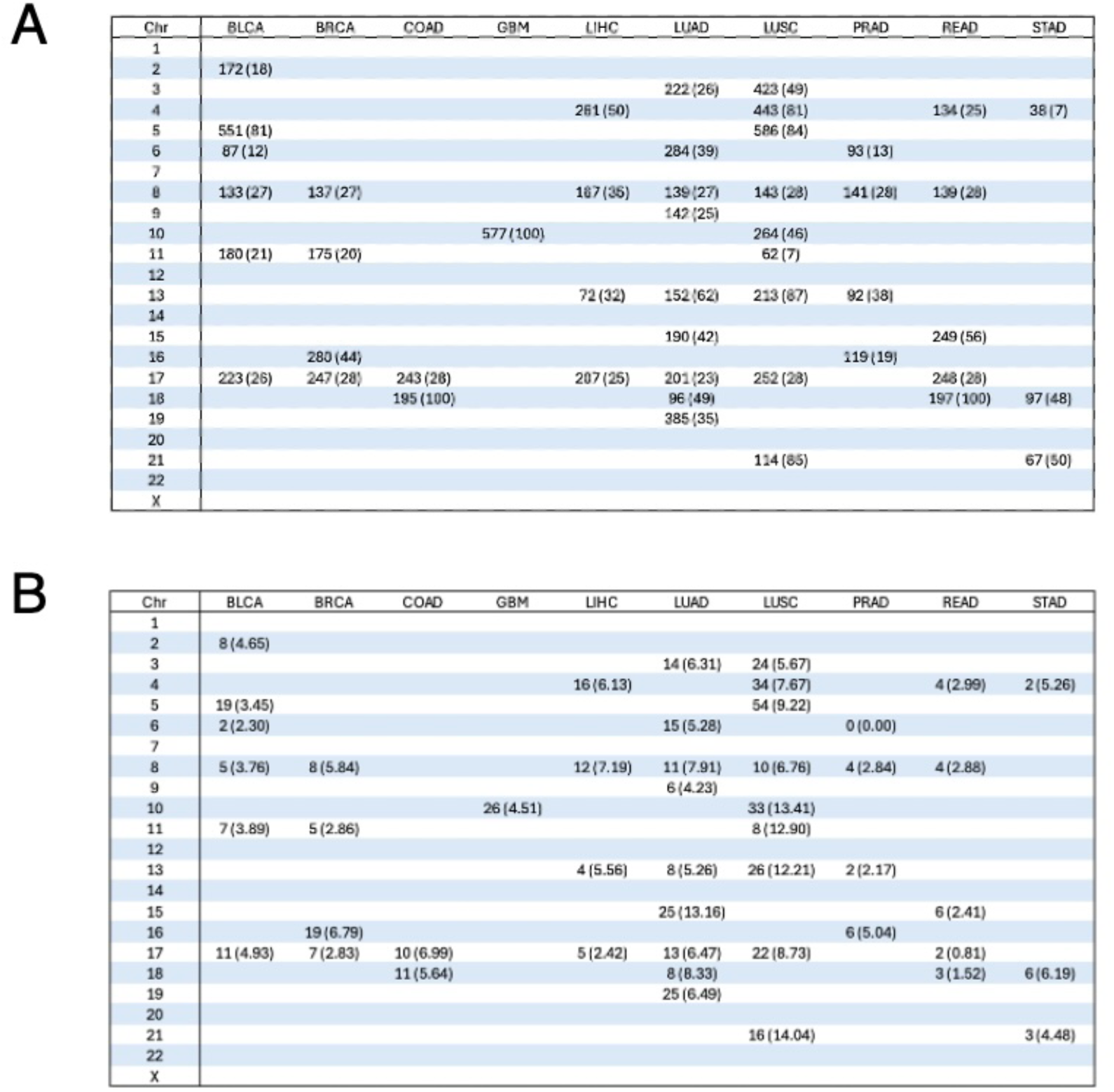
Genome-wide analysis of regions with recurrent copy number loss. A) For all chromosomes across ten cancer datasets, the number of genes within recurrently deleted regions is shown. The percentage of genes on the chromosome present in the deleted region is shown in parentheses. Only genes that are expressed above an average log2 cut-off of 2 in either normal or tumor in the dataset are included. B) For each of the recurrent copy number loss regions shown in A, the number of genes with higher expression in tumor than normal (T-N log2FC > 1, p-adj < 0.0001) is shown. The number in parentheses indicates the percentage of expressed genes within the region that meet this differential expression cut-off.

**Figure S4.**
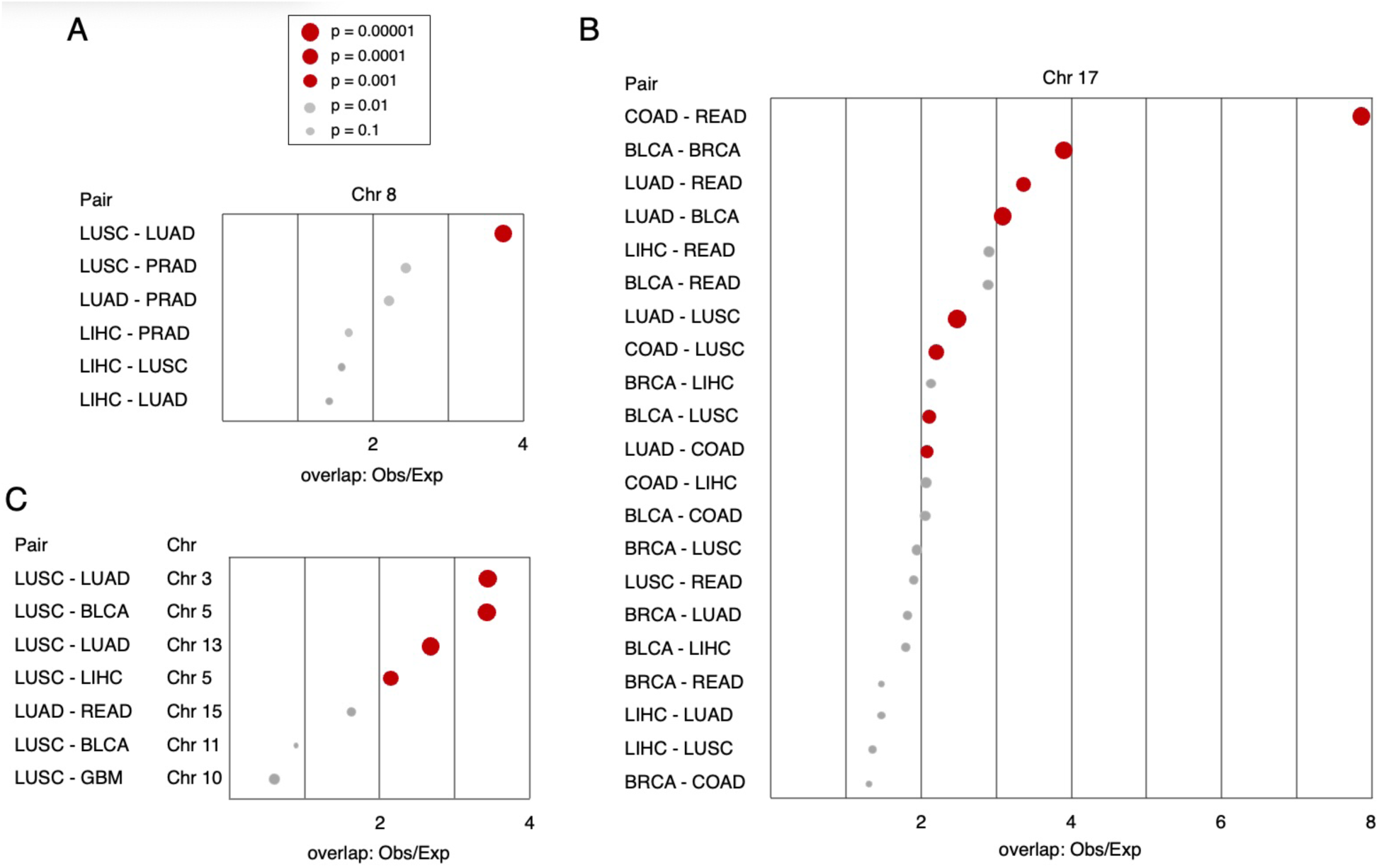
Analysis of pairwise overlap between upregulated genes among cancer types. For all pairs of cancers with a shared copy number reduced region where there were at least 15 genes with higher expression in tumor than normal (at the more relaxed cutoff: T-N log2FC > 0.4, p-adj < 0.01), the significance of the overlap was analyzed (hypergeometric test). A and B show all pairwise comparisons for the regions on chromosome 8 (A) and chromosome 17 (B). Panel C shows a similar analysis for all other regions with at least 15 genes upregulated in each of the indicated pairs. Cancer pairs are shown on the left (the chromosomal region is also shown for C). Data is plotted as fold difference comparing the observed overlap to the expected, with the size of each dot representing the p-value. The size key for all p-values is shown in A: p < 0.01 shown in red, p >= 0.01 in gray.

**Figure S5.**
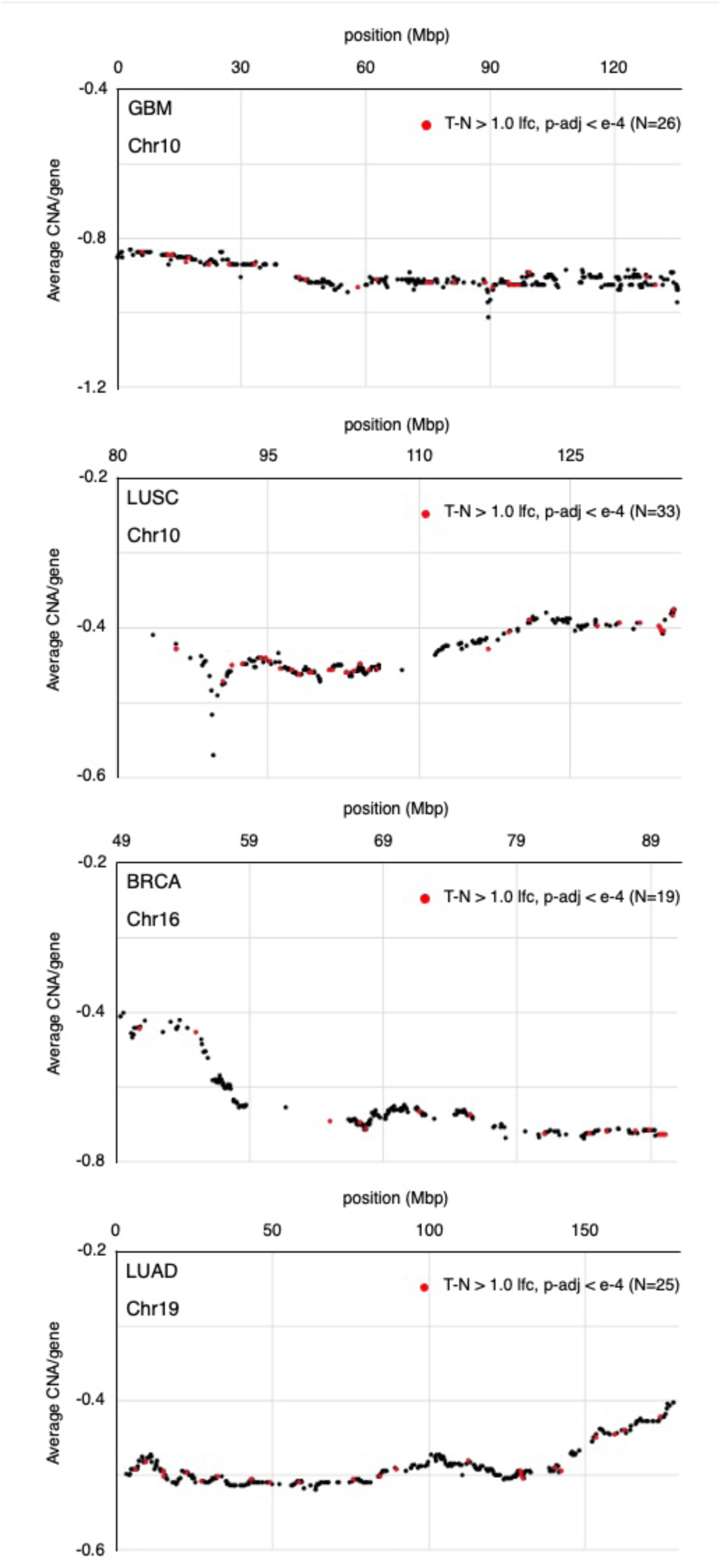
Genes with higher expression in tumor than normal are scattered across deletion regions. Average copy number for all expressed genes within recurrent copy number loss regions on chromosome 10 (GBM and LUSC), chromosome 16 (BRCA) and chromosome 19 (LUAD) is plotted. The positions of genes with higher expression in tumor than normal (T-N log2FC > 1, p-adj < 0.0001) is shown in red. The number of genes meeting the differential expression cut-off is shown for each plot. Note that different amounts of chromosome 10 are plotted for GBM and LUSC, since the recurrent copy number reduction is different in these two datasets, encompassing 100% of the chromosome in GBM but only ∼46% in LUSC. These four regions were selected since they were relatively large and included higher numbers of genes with higher expression in the tumor samples.

**Figure S6.**
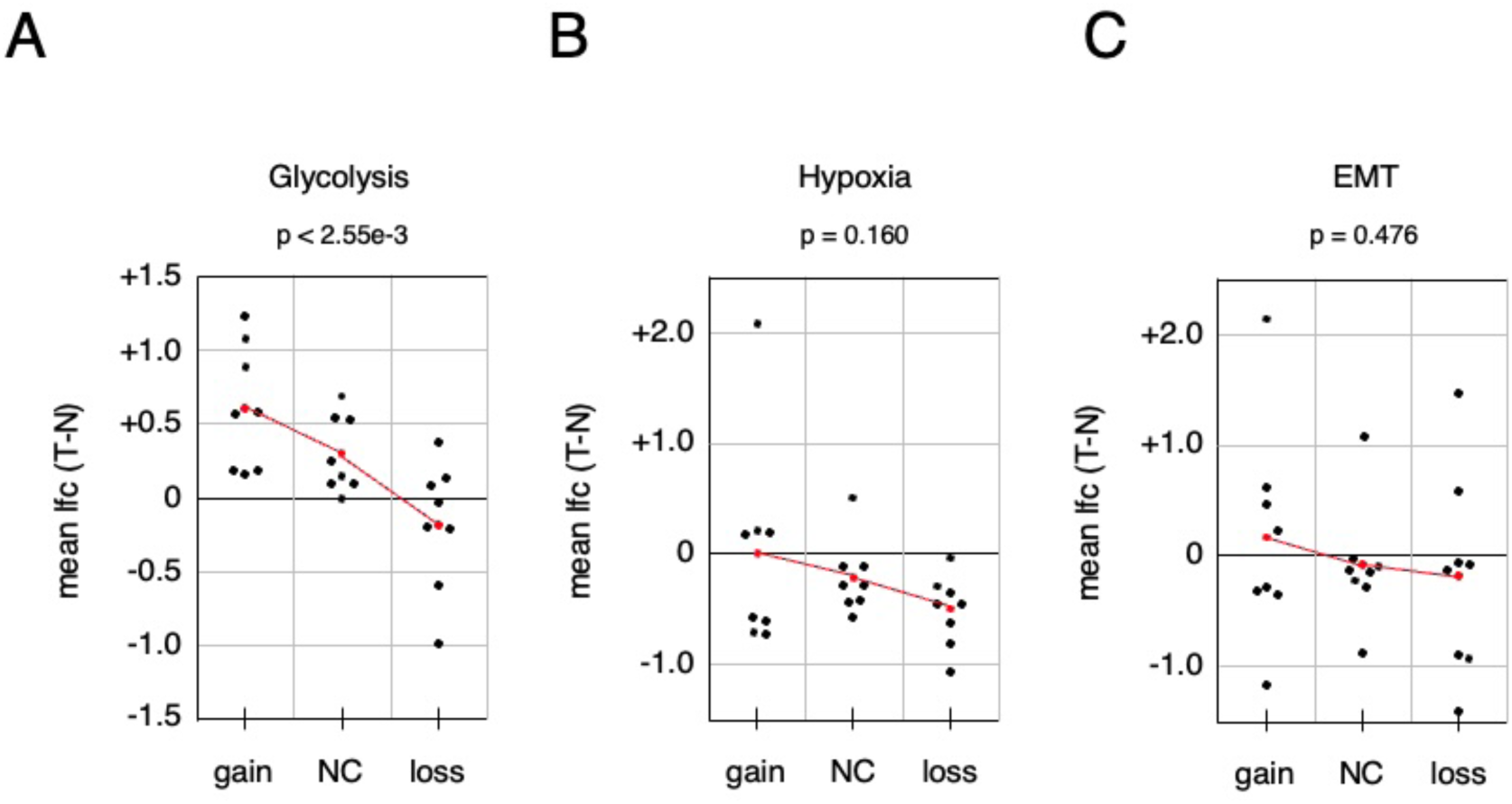
Gene expression changes and copy number variation. Each dataset was divided by copy number (loss: < -0.4, gain: > +0.4, NC: -0.4 to +0.4) and average T-N log2 gene expression difference was plotted for each of the indicated gene sets: A) Glycolysis, B) Hypoxia, C) EMT. p-values are for the comparison of average log2FC between the gain and loss categories. Plotted in red is the average for each bin and the trendline based on the average.

## References

1. Alexandrov LB, Nik-Zainal S, Wedge DC, Aparicio SAJR, Behjati S, Biankin AV, et al. Signatures of mutational processes in human cancer. Nature. 2013;500: 415–421. doi:10.1038/nature12477

2. Zack TI, Schumacher SE, Carter SL, Cherniack AD, Saksena G, Tabak B, et al. Pan-cancer patterns of somatic copy number alteration. Nat Genet. 2013;45: 1134–1140. doi:10.1038/ng.2760

3. Beroukhim R, Mermel CH, Porter D, Wei G, Raychaudhuri S, Donovan J, et al. The landscape of somatic copy-number alteration across human cancers. Nature. 2010;463: 899–905. doi:10.1038/nature08822

4. Davoli T, Xu AW, Mengwasser KE, Sack LM, Yoon JC, Park PJ, et al. Cumulative haploinsufficiency and triplosensitivity drive aneuploidy patterns and shape the cancer genome. Cell. 2013;155: 948–962. doi:10.1016/j.cell.2013.10.011

5. Davies H, Bignell GR, Cox C, Philip Stephens, Edkins S, Clegg S, et al. Mutations of the BRAF gene in human cancer. Nature. 2002;417: 949–954. doi:10.1038/nature00766

6. Simanshu DK, Nissley DV, McCormick F. RAS Proteins and Their Regulators in Human Disease. Cell. 2017;170: 17–33. doi:10.1016/j.cell.2017.06.009

7. Chen X, Zhang T, Su W, Dou Z, Zhao D, Jin X, et al. Mutant p53 in cancer: from molecular mechanism to therapeutic modulation. Cell Death Dis. 2022;13: 974. doi:10.1038/s41419-022-05408-1

8. Fearon ER. Molecular genetics of colorectal cancer. Annual review of pathology. 2011;6: 479–507.

9. Sparks AB, Morin PJ, Vogelstein B, Kinzler KW. Mutational analysis of the APC/beta-catenin/Tcf pathway in colorectal cancer. Cancer Res. 1998;58: 1130–1134.

10. Smith G, Carey FA, Beattie J, Wilkie MJV, Lightfoot TJ, Coxhead J, et al. Mutations in APC, Kirsten-ras, and p53--alternative genetic pathways to colorectal cancer. Proc Natl Acad Sci U S A. 2002;99: 9433–9438. doi:10.1073/pnas.122612899

11. Wang SI, Puc J, Li J, Bruce JN, Cairns P, Sidransky D, et al. Somatic mutations of PTEN in glioblastoma multiforme. Cancer Res. 1997;57: 4183–4186.

12. Tarafa G, Villanueva A, Farré L, Rodríguez J, Musulén E, Reyes G, et al. DCC and SMAD4 alterations in human colorectal and pancreatic tumor dissemination. Oncogene. 2000;19: 546–555. doi:10.1038/sj.onc.1203353

13. Kinzler KW, Vogelstein B. Lessons from hereditary colorectal cancer. Cell. 1996;87: 159–70.

14. Hanahan D. Hallmarks of Cancer: New Dimensions. Cancer Discov. 2022;12: 31–46. doi:10.1158/2159-8290.CD-21-1059

15. Hanahan D, Weinberg RA. The hallmarks of cancer. Cell. 2000;100: 57–70.

16. Gordon DJ, Resio B, Pellman D. Causes and consequences of aneuploidy in cancer. Nat Rev Genet. 2012;13: 189–203. doi:10.1038/nrg3123

17. Ben-David U, Amon A. Context is everything: aneuploidy in cancer. Nat Rev Genet. 2020;21: 44–62. doi:10.1038/s41576-019-0171-x

18. Bertolino E, Reimund B, Wildt-Perinic D, Clerc R. A novel homeobox protein which recognizes a TGT core and functionally interferes with a retinoid-responsive motif. J Biol Chem. 1995;270: 31178–31188.

19. Wotton D, Lo RS, Lee S, Massague J. A Smad transcriptional corepressor. Cell. 1999;97: 29–39.

20. Melhuish TA, Gallo CM, Wotton D. TGIF2 interacts with histone deacetylase 1 and represses transcription. J Biol Chem. 2001;276: 32109–32114.

21. Massague J. TGFbeta in Cancer. Cell. 2008;134: 215–30.

22. Shah A, Melhuish TA, Fox TE, Frierson HF, Wotton D. TGIF transcription factors repress acetyl CoA metabolic gene expression and promote intestinal tumor growth. Genes Dev. 2019;33: 388–402. doi:10.1101/gad.320127.118

23. Kong L, Yu Y, Guan H, Jiang L, Sun F, Li X, et al. TGIF1 plays a carcinogenic role in esophageal squamous cell carcinoma through the Wnt/β-catenin and Akt/mTOR signaling pathways. Int J Mol Med. 2021;47: 77. doi:10.3892/ijmm.2021.4910

24. Wang Y, Pan T, Li L, Wang H, Li J, Zhang D, et al. Knockdown of TGIF attenuates the proliferation and tumorigenicity of EC109 cells and promotes cisplatin-induced apoptosis. Oncol Lett. 2017;14: 6519–6524. doi:10.3892/ol.2017.7009

25. Wang B, Ma Q, Wang X, Guo K, Liu Z, Li G. TGIF1 overexpression promotes glioma progression and worsens patient prognosis. Cancer Med. 2022;11: 5113–5128. doi:10.1002/cam4.4822

26. Zhang T, Song X, Zhang Z, Mao Q, Xia W, Xu L, et al. Aberrant super-enhancer landscape reveals core transcriptional regulatory circuitry in lung adenocarcinoma. Oncogenesis. 2020;9: 92. doi:10.1038/s41389-020-00277-9

27. Parajuli P, Singh P, Wang Z, Li L, Eragamreddi S, Ozkan S, et al. TGIF1 functions as a tumor suppressor in pancreatic ductal adenocarcinoma. Embo J. 2019. doi:10.15252/embj.2018101067

28. Weng CC, Hsieh MJ, Wu CC, Lin YC, Shan YS, Hung WC, et al. Loss of the transcriptional repressor TGIF1 results in enhanced Kras-driven development of pancreatic cancer. Mol Cancer. 2019;18: 96. doi:10.1186/s12943-019-1023-1

29. Cornish AJ, Gruber AJ, Kinnersley B, Chubb D, Frangou A, Caravagna G, et al. The genomic landscape of 2,023 colorectal cancers. Nature. 2024;633: 127–136. doi:10.1038/s41586-024-07747-9

30. Nunes L, Li F, Wu M, Luo T, Hammarström K, Torell E, et al. Prognostic genome and transcriptome signatures in colorectal cancers. Nature. 2024;633: 137–146. doi:10.1038/s41586-024-07769-3

31. Chen GT, Tifrea DF, Murad R, Habowski AN, Lyou Y, Duong MR, et al. Disruption of β-Catenin-Dependent Wnt Signaling in Colon Cancer Cells Remodels the Microenvironment to Promote Tumor Invasion. Mol Cancer Res. 2022;20: 468–484. doi:10.1158/1541-7786.MCR-21-0349

32. Melhuish TA, Adair SJ, Pemberton OS, Bauer TW, Wotton D. A simple method for analyzing competitive growth of multiple cell types in xenograft tumors. bioRxiv. 2026; 2026.01.23.701386. doi:10.64898/2026.01.23.701386

33. Arafeh R, Shibue T, Dempster JM, Hahn WC, Vazquez F. The present and future of the Cancer Dependency Map. Nat Rev Cancer. 2025;25: 59–73. doi:10.1038/s41568-024-00763-x

34. Sadasivam S, Duan S, DeCaprio JA. The MuvB complex sequentially recruits B-Myb and FoxM1 to promote mitotic gene expression. Genes Dev. 2012;26: 474–489. doi:10.1101/gad.181933.111

35. Madison BB, Dunbar L, Qiao XT, Braunstein K, Braunstein E, Gumucio DL. Cis elements of the villin gene control expression in restricted domains of the vertical (crypt) and horizontal (duodenum, cecum) axes of the intestine. J Biol Chem. 2002;277: 33275–83. doi:10.1074/jbc.M204935200

36. Hahn SA, Schutte M, Hoque ATMS, Moskaluk CA, da Costa LT, Rozenblum E, et al. DPC4, a candidate tumor suppressor gene at human chromosome 18q21.1. Science. 1996;271: 350–353.

37. Levy L, Hill CS. Alterations in components of the TGF-beta superfamily signaling pathways in human cancer. Cytokine & growth factor reviews. 2006;17: 41–58.

38. Takaku K, Oshima M, Miyoshi H, Matsui M, Seldin MF, Taketo MM. Instestinal tumorigenesis in compound mutant mice of both *Dpc4 (Smad4)* and *APC* genes. Cell. 1998;92: 645–656.

39. Chen S, Francioli LC, Goodrich JK, Collins RL, Kanai M, Wang Q, et al. A genomic mutational constraint map using variation in 76,156 human genomes. Nature. 2024;625: 92–100. doi:10.1038/s41586-023-06045-0

40. Wotton D, Taniguchi K. Functions of TGIF homeodomain proteins and their roles in normal brain development and holoprosencephaly. American journal of medical genetics. 2018. doi:10.1002/ajmg.c.31612

41. Pfister K, Pipka JL, Chiang C, Liu Y, Clark RA, Keller R, et al. Identification of Drivers of Aneuploidy in Breast Tumors. Cell Rep. 2018;23: 2758–2769. doi:10.1016/j.celrep.2018.04.102

42. Laoukili J, Kooistra MRH, Brás A, Kauw J, Kerkhoven RM, Morrison A, et al. FoxM1 is required for execution of the mitotic programme and chromosome stability. Nat Cell Biol. 2005;7: 126–136. doi:10.1038/ncb1217

43. Chien JC-Y, Tabet E, Pinkham K, da Hora CC, Chang JC-Y, Lin S, et al. A multiplexed bioluminescent reporter for sensitive and non-invasive tracking of DNA double strand break repair dynamics in vitro and in vivo. Nucleic Acids Res. 2020;48: e100. doi:10.1093/nar/gkaa669

44. Melhuish TA, Kowalczyk I, Manukyan A, Zhang Y, Shah A, Abounader R, et al. Myt1 and Myt1l transcription factors limit proliferation in GBM cells by repressing YAP1 expression. Biochim Biophys Acta Gene Regul Mech. 2018;1861: 983–995. doi:10.1016/j.bbagrm.2018.10.005

45. Truett GE, Heeger P, Mynatt RL, Truett AA, Walker JA, Warman ML. Preparation of PCR-quality mouse genomic DNA with hot sodium hydroxide and tris (HotSHOT). Biotechniques. 2000;29: 52, 54.

46. Cerami E, Gao J, Dogrusoz U, Gross BE, Sumer SO, Aksoy BA, et al. The cBio cancer genomics portal: an open platform for exploring multidimensional cancer genomics data. Cancer Discov. 2012;2: 401–404. doi:10.1158/2159-8290.CD-12-0095

47. Chen EY, Tan CM, Kou Y, Duan Q, Wang Z, Meirelles GV, et al. Enrichr: interactive and collaborative HTML5 gene list enrichment analysis tool. BMC Bioinformatics. 2013;14: 128. doi:10.1186/1471-2105-14-128

48. Kuleshov MV, Jones MR, Rouillard AD, Fernandez NF, Duan Q, Wang Z, et al. Enrichr: a comprehensive gene set enrichment analysis web server 2016 update. Nucleic Acids Res. 2016;44: W90–7. doi:10.1093/nar/gkw377

49. Goldman MJ, Craft B, Hastie M, Repečka K, McDade F, Kamath A, et al. Visualizing and interpreting cancer genomics data via the Xena platform. Nat Biotechnol. 2020;38: 675–678. doi:10.1038/s41587-020-0546-8

